# YAP1-induced super-enhancer rewiring mediates gene–environment-driven malignant reprogramming and metastatic ecosystem formation in head and neck cancer

**DOI:** 10.64898/2026.11.15.766730

**Authors:** Masaaki Higashihara, Kuniaki Sato, Hirofumi Omori, Noritaka Komune, Takahiro Hongo, Takuya Nakagawa, Masashige Bando, Ryuichiro Nakato, Fumihito Miura, Hiromitsu Araki, Takafumi Nakano, Kenichi Taguchi, Takashi Nakagawa, Takashi Ito, Katsuhiko Shirahige, J Silvio Gutkind, Atsushi Kaneda, Muneyuki Masuda

## Abstract

Head and neck squamous cell carcinoma (HNSCC) is characterized by frequent *TP53* mutations, which are insufficient for tumorigenesis and largely undruggable, underscoring the need to define epigenetic mechanisms driving cancer evolution. Here we show that *TP53* mutation facilitates YAP1-dependent transcriptional reprogramming, driving epigenetic tumor evolution by hijacking tissue-regenerative programs. Integrated multi-omics analyses of mouse models and multiregional human HNSCC samples reveal that global chromatin architecture remains largely stable during progression, whereas YAP1 dynamically redistributes to newly formed super-enhancers accompanied by enhancer DNA hypomethylation. Mechanistically, YAP1 establishes an IL6-dependent feedforward loop and cooperates with stage-specific transcription factors, including FOXM1, ΔNp63, and PITX2. The YAP1–PITX2 axis promotes partial EMT, cancer-associated fibroblast activation, and lymph node metastasis, resulting in a fibrotic and immunosuppressive metastatic ecosystem. These findings identify a YAP1-driven epigenetic circuit as a central hub linking environmental stress to malignant reprogramming and highlight a potential therapeutic vulnerability in HNSCC.

## Introduction

Head and neck squamous cell carcinoma (HNSCC) is the sixth most common cancer in the world, affecting 600,000 people annually. Despite favorable clinical outcomes in patients with Human papillomavirus (HPV)-positive HNSCC, both the quantity and quality of life of patients with aggressive HPV-negative HNSCC, have not improved over the past decades: approximately half of patients die from their disease, while survivors frequently suffer substantial functional impairment due to the loss of critical organs (e.g., the larynx) ^1,2^. Hereafter, unless otherwise specified, HNSCC refers to the HPV-negative HNSCC. Moreover, precision medicine has shown limited success in HNSCC, because this cancer is characterized by frequent mutations in unactionable tumor suppressor gene (TSG), including *TP53* (84%), *CDKN2A* (57%), *FAT1*(32%), and *NOTCH 1(*26%) ^3,4^. In addition to this clinical conundrum, genome-oriented views fail to fully account for the biology of HNSCC. Recent evidence indicates that these mutations accumulate prior to malignant transformation in the normal epithelium of the upper-aerodigestive-tract, and that their composition remains largely unchanged when tumors recur or metastasize to lymph nodes ^5-7^. In contrast, cancer cells are distinguished from normal cells by dysregulated epigenetic states that can be acquired in the presence of gene mutations ^8-10^. These epigenetic alterations enable cancer cells to override homeostatic regulatory signals and flexibly adapt to environmental stress through remodeling of the transcriptional machinery. For example, the formation and hypomethylation of super-enhancers (SEs) promote malignant reprogramming by converting environmental oncogenic inputs into amplified transcriptional outputs ^11-13^. Accordingly, cancer has been proposed to represent a gene-environment-driven epigenetic process, often described as transcriptional addiction ^8,9,11,14^. This concept aligns with the field carcinogenesis theory, which has long been applied to explain HNSCC^15^. HNSCC-associated environmental stimuli—such as tobacco, alcohol, and chronic inflammation— predominantly induce TSG mutations while simultaneously driving cellular-context-dependent oncogenic transcriptional programs, including tissue regeneration, matrix remodeling, de-differentiation, epithelial-mesenchymal transition (EMT), acquisition of stemness, and immune evasion ^16,17^. We, therefore, hypothesized that HNSCC represents an adaptive system that is highly dependent on gene-environment-driven transcriptional reprograming ^2,18^.

In this context, we previously investigated the roles of transcriptional coactivator YAP1 in HNSCC ^19,20^. YAP1 functions as a sensor of microenvironment and transmits biological cues to regenerative transcriptional programs in epithelial stem/progenitor cells ^21^. Endogenous YAP1 activity is essential for maintaining the rapid turn-over of the squamous epithelium (e.g., approximately 14 days in the oral mucosa) and for promoting tissue regeneration under acute stress conditions such as tissue injury ^21^. In our previous study, conditional activation of YAP1 induced ultra-rapid tumor formation in the tongue of mice: carcinoma *in situ* (CIS) within 2 weeks and invasive SCC within 4 weeks, with 100% reproducibility ^19^. In clinical HNSCC specimens, elevated YAP1 activity was significantly associated with lymph node (LN) metastasis, tumor recurrence, and unfavorable prognosis^19^. In a recent study, we further demonstrated that YAP1 signaling drives malignant reprograming of external auditory canal SCC, a rare subtype of HNSCC mainly caused by chronic inflammation^22^. These findings indicate that excessive activation of endogenous tissue-repair transcriptional programs can drive tumorigenesis, recapitulating the concept of “wounds that do not heal” ^23^. Furthermore, we demonstrated that YAP1, triggered by environmental stressors, transcriptionally reprograms HNSCC cells toward the acquisition of stemness ^20^.

Beyond our study, the potent oncogenic roles of YAP1 have been validated across a wide spectrum of malignancies including skin, liver, breast, stomach, pancreas, and brain ^24-27^. Mechanistically, YAP1 acts as an enhancer-binding transcriptional coactivator that promotes the assembly—via phase separation—of SEs together with p300, MED1, BRD4 and partner transcription factors (TFs) —such as TEAD, AP-1, and Smad—thereby amplifying transcription of target genes through the recruitment of RNA polymerase II ^25,28-34^. Canonical regulation of YAP1 activity has been attributed primarily to negative control by the Hippo pathway through serine127 phosphorylation, leading to cytoplasmic sequestration and proteasomal degradation ^35^. However, an increasing body of evidence demonstrates that multiple tyrosine kinases—including Src, FAK, and c-Abl—can activate YAP via phosphorylation at tyrosine (Y) 357 ^36-38^. Notably, these kinases are activated by extracellular cues that are abundant in the HNSCC-specific tumor microenvironment (TME), including IL6, collagen, integrin, laminin, and TGF-b and its latent complex deposited in the extracellular matrix ^39,40^. Moreover, YAP1 activity is required for the maintenance of TGF-b-induced cancer-associated fibroblasts (CAFs), which serve as key mediators of matrix remodeling and tissue fibrosis that reciprocally reinforce YAP1 activation in both CAFs and cancer cells ^17,41^. In parallel, emerging evidence suggests that p53 and FAT1 may modulate the Hippo pathway and thereby influence YAP1 signaling. In this context, the high prevalence of *TP53* and *FAT1* mutations in HNSCC may contribute to YAP1-driven cellular reprogramming ^42-44^. Collectively, these findings support that YAP1 functions as a central communication hub linking HNSCC cells to their TME.

Building on these observations, we hypothesized that tumor evolution in HNSCC is driven primarily by gene–environment–induced transcriptional reprogramming rather than by additional genetic alterations. In particular, we postulated that *TP53* mutation facilitates YAP1 activation, enabling cancer cells to convert microenvironmental signals into oncogenic transcriptional programs through enhancer remodeling. To test this hypothesis, we combined epigenetic carcinogenesis models, multi-omics analyses of multiregional human HNSCC samples, and spatial transcriptomic profiling of tumor microenvironments. Our results reveal that global chromatin architecture remains largely stable during HNSCC progression, whereas YAP1 dynamically redistributes to newly formed super-enhancers accompanied by enhancer DNA hypomethylation. Notably, *TP53* mutation facilitates this process by relieving p53-mediated constraint on YAP1, thereby enabling sustained YAP1 activity. This selective enhancer rewiring establishes stage-specific transcriptional programs that promote tumor progression, lymph node metastasis, and formation of a fibrotic and immunosuppressive tumor microenvironment.

## Results

### Clinical significance of YAP1-induced transcriptional program and its modulation by *TP53* mutation

In our initial study, we examined the clinical significance of YAP1-induced transcriptional program. Employing a cell line derived from the tamoxifen (*TAM*)-inducible *Mob1a/b* double-knock-out (DKO) mouse [*Rosa26-CreERT*; *Mob1a^flox/flox^*; *Mob1b^-/-^* (tg*Mob1*DKO) mice](*iMob1*DKO cell), which we established in our previous study ^19^, the gene expression profile was analyzed at the timing of dysplastic change, i.e., 1 w after TAM treatment (Fig.1a, and Supplementary Data 1). Gene set enrichment assays (GSEA) showed that up-regulated genes (Z score >2) revealed significant enrichment of regenerative and proliferative programs, including E2F targets, G2M checkpoint, mitotic spindle, and EMT (Fig. 1b), which was further supported by complementary individual GSEA enrichment plots and Hallmark enrichment analysis (Supplementary Fig. 1a, b). Of note, the p53 pathway showed concurrent negative enrichment in this YAP1-driven regenerative state (Fig.1b), supporting a model in which antagonistic p53–YAP1 signaling governs cellular plasticity ^43^, whereby p53 constrains dedifferentiation-associated reprogramming while YAP1 promotes regenerative transcriptional remodeling. We then selected the top 16 up-regulated genes in reference to the public YAP1 target gene data ^31^: *TK1, DAB2, BIRC5, TOP2A, CENPF, FSCN1, IGFBP3, SHCBP1, TNNT2, CDC20, THBS1, ANKRD1, STMN1, HMMR, AXL,* and *CTGF,* and termed them as a YAP1 carcinogenesis module (YCM) (Fig.1c). In the human TCGA data of HNSCC patients with high YCM activity was associated with significantly poor prognosis (*P* = 0.0093) (Fig.1d). We re-examined the prognostic value of YAP1 protein expression in the HNSCC clinical samples obtained at National Kyushu Cancer Center expanding the number of patients from 86 (analyzed in the previous study^19^) to 119 and updating their clinical data (Table 1). As shown in the representative case, endogenous YAP1 expression spreads according to the tumor progression (i.e., normal, dysplasia, SCC and invasive SCC) (Fig.1e) and worsened the overall survival of patients (*P* = 0.012, log-rank test) (Fig.1f). In the multivariate analysis of clinicopathological factors, YAP1 expression (HR:2.54; 95% CI:1.13-5.68; *P* = 0.024) and age were independent prognostic factors irrespective of tumor stage (Table 1).

**Figure 1.**
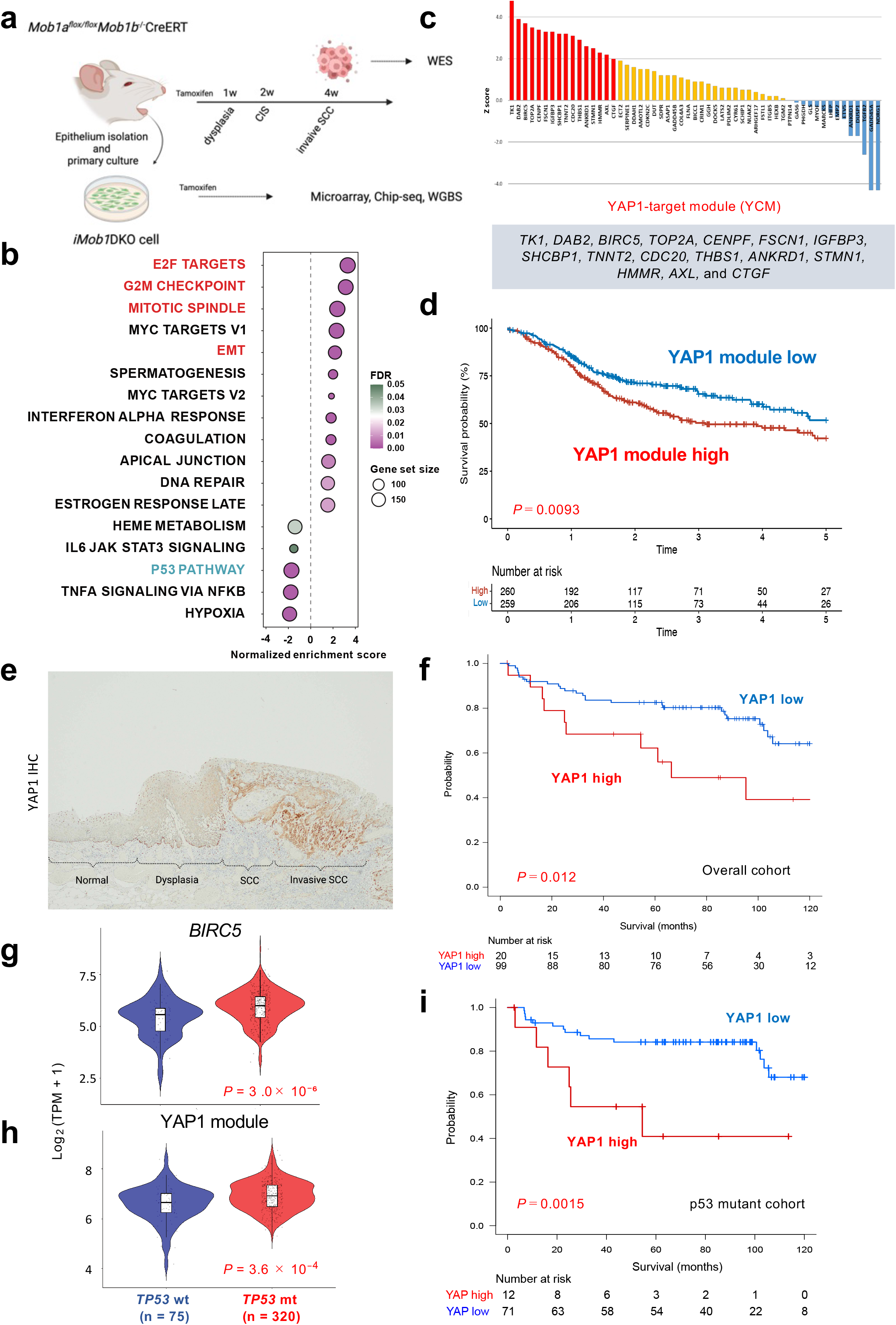
Clinical significance of the YAP1-induced transcriptional program and its modulation by *TP53* mutation. **a** Experimental design illustrating tumor development and cell line derivation in a tamoxifen (TAM)-inducible *Mob1a/b* double-knockout (*iMob1*DKO) mouse model, followed by multi-omics profiling of tumor tissues and derived cells. Created in BioRender. https://BioRender.com/BL29Z4ZTY4. **b** Hallmark gene sets enriched in tongue epithelial tissues of tamoxifen-induced *Mob1* double-knockout (DKO) mice at 1 week after tamoxifen treatment. The x-axis represents the normalized enrichment score (NES), circle size indicates gene set size, and color indicates the false discovery rate (FDR) q value. **c** Bar plot showing Z score ranking of genes altered in *iMob1*DKO cells following YAP1 activation. Integration with publicly available YAP1 target gene datasets identified 16 upregulated genes (Z score > 2), which were defined as the YAP1 carcinogenesis module (YCM). **d** Kaplan–Meier analysis of overall survival in TCGA HNSCC patients stratified by YAP1 carcinogenesis module (YCM) activity according to single sample GSEA (ssGSEA) scores, showing poorer prognosis in patients with high YCM activity (*P* = 0.0093, log-rank test). **e** Representative immunohistochemical staining showing progressive expansion of endogenous YAP1 expression across histological stages (normal epithelium, dysplasia, SCC, and invasive SCC). **f** Kaplan–Meier analysis of overall survival in HNSCC clinical samples stratified by nuclear YAP1 protein expression (n = 119), showing significantly worse survival in YAP1-high tumors (*P* = 0.012, log-rank test). **g, h** *TP53* mutation is associated with activation of the YAP1 transcriptional program. Violin plots showing increased expression of BIRC5 (g) and elevated YCM activity (h) in *TP53*-mutant tumors compared with *TP53* wild-type tumors (*P* = 3 × 10□□ and *P* = 3.6 × 10□□, respectively). P values were calculated using a two-sided Wilcoxon rank-sum test. **i** Kaplan–Meier analysis of overall survival in *TP53*-mutant HNSCC clinical samples stratified by nuclear YAP1 protein expression (n = 83), showing significantly worse survival in YAP1-high tumors (*P* = 0.0015, log-rank test). See Source Data for **b, f, g, h** and **i.**

**Table 1.** Univariable and multivariable Cox proportional hazards analyses of clinicopathological factors associated with overall survival in patients with HNSCC.

| **Variable** | **All** |  | **Comparison** | **Univariable analysis** |  |  | **Multivariable analysis** |  |  |
|---|---|---|---|---|---|---|---|---|---|
|  | **n = 119** | **%** |  | **HR** | **95% CI** | **P** | **HR** | **95% CI** | **P** |
| **Age** |  |  |  |  |  |  |  |  |  |
| Median (range) | 64 (22–87) |  | ≥64/<64 | 2.35 | 1.19–4.65 | 0.014 | 2.53 | 1.26–5.08 | 0.009 |
| ≥64 years | 60 | 50.4 |  |  |  |  |  |  |  |
| <64 years | 59 | 49.6 |  |  |  |  |  |  |  |
| **Sex** |  |  |  |  |  |  |  |  |  |
| Male | 85 | 71.4 | Male/Female | 1.70 | 0.75–3.88 | 0.204 |  |  |  |
| Female | 34 | 28.6 |  |  |  |  |  |  |  |
| **T stage** |  |  |  |  |  |  |  |  |  |
| Low (T1–T2) | 88 | 73.9 | High/Low | 1.27 | 0.63–2.55 | 0.512 |  |  |  |
| High (T3–T4) | 31 | 26.1 |  |  |  |  |  |  |  |
| **N stage** |  |  |  |  |  |  |  |  |  |
| N0 | 92 | 77.3 | N1–3/N0 | 1.46 | 0.71–3.00 | 0.303 |  |  |  |
| N1–3 | 27 | 22.7 |  |  |  |  |  |  |  |
| **Differentiation** |  |  |  |  |  |  |  |  |  |
| Well to moderate | 110 | 92.4 | Poor/Well–moderate | 2.40 | 1.01–5.77 | 0.049 | 1.26 | 0.46–3.43 | 0.655 |
| Poor | 9 | 7.6 |  |  |  |  |  |  |  |
| **YAP1 expression** |  |  |  |  |  |  |  |  |  |
| Low | 99 | 83.2 | High/Low | 2.40 | 1.19–4.83 | 0.014 | 2.54 | 1.13–5.68 | 0.024 |
| High | 20 | 16.8 |  |  |  |  |  |  |  |

HR, hazard ratio; CI, confidence interval; HNSCC, head and neck squamous cell carcinoma. P values were derived from Cox proportional hazards regression analyses.

We then analyzed whether this YAP1-induced cellular reprograming is relevant to the transcriptional level of YAP1. In the TCGA data, the levels of YAP1 mRNA were similar between normal and tumor samples (*P* = 0.85) (Supplementary Fig.1c) and demonstrated no survival significance (*P* = 0.2) (Supplementary Fig.1d) in HNSCC patients, contrary to the nuclear YAP1 protein level and YCM activity (Fig.1d, f and Table 1). Collectively, post-translational modification of YAP1 protein and YAP1-induced transcriptional program appears to be a *bona fide* driver of human HNSCC.

In light of the proposed antagonistic p53–YAP1 signaling axis and the established inhibitory role of wild-type p53 in YAP1-driven carcinogenesis in squamous epithelium ^42^, we examined whether the frequent *TP53* mutations observed in HNSCC facilitate YAP1-induced cellular reprogramming. Analysis of TCGA datasets revealed that *TP53* mutation was significantly associated with enhanced activity of YAP1-target-genes, *BIRC5* (*P* = 3 × 10^-6^) and YCM (*P* = 3.6 × 10^-4^), (Fig. 1g, h), whereas *YAP1* mRNA levels were not affected (Supplementary Fig.1e). We next examined p53 protein status by immunohistochemistry in the same human HNSCC cohort analyzed for YAP1 expression (Supplementary Fig.1f). In the *TP53*-mutant subgroup (n = 83), the prognostic impact of YAP1 overexpression was more prominent (*P* = 0.0015; Fig.1i) than in the overall cohort (n = 119) (Fig.1f). Notably, among the 8 cases, in which mutant p53 expression was detected in cancer-adjacent normal epithelium, frequent (68.5%) colocalization of YAP1 was observed (Supplementary Fig.1e). These findings suggest that gene-environment interactions between *TP53* mutation and YAP1 activation may be initiated at a pre-carcinogenic stage. Taken together with our previous findings that *TP*53 knockout or mutation enhances nuclear retention of YAP1 in HNSCC cell lines and clinical samples ^19^, these results indicate that *TP*53 mutations—present in the majority of HNSCC cases (84%)—potentiate YAP1-induced malignant reprograming predominantly through post-translational mechanisms.

### Effects of YAP1 on global chromatin conformation in a mouse carcinogenesis model

Given the clinical significance of YAP1-driven transcriptional programs in HNSCC, we sought to elucidate the epigenetic mechanisms underlying YAP1-induced carcinogenic reprogramming using an inducible *Mob1* double-knockout (*iMob1*DKO) mouse model (Fig. 1a). Prior to epigenetic profiling, we assessed the genetic landscape of pathologically confirmed invasive carcinomas that developed in mice 4 weeks after tamoxifen (TAM) treatment by whole-exome sequencing (Fig. 1a). We used the Gene database of the National Center for Biotechnology Information (NCBI) to assess the genes harboring somatic variants and found none that we classified as cancer-associated. This observation raised the possibility that epigenetic changes contribute to YAP1-driven carcinogenesis in this mouse model. We therefore proceeded with epigenetic analyses, including chromatin immunoprecipitation sequencing (ChIP-seq) and whole-genome bisulfite sequencing (WGBS), using *iMob1*DKO cells collected at 1 and 2 weeks after TAM treatment, corresponding to the stages of dysplasia and carcinoma in situ (CIS), respectively (Fig. 1a). Given the rapid progression of epigenetic carcinogenesis in this model, we anticipated substantial alterations in facultative heterochromatin marked by H3K9me2 and in enhancer activity marked by H3K27ac. However, the mutually exclusive distribution and relative intensities of H3K9me2 and H3K27ac remained largely unchanged across both broad genomic regions and narrow transcription start site (TSS)–proximal domains following TAM treatment (Supplementary Fig. 2a, b). In contrast, WGBS revealed pronounced alterations in DNA methylation patterns. YAP1 activation markedly increased the number of differentially methylated regions (DMRs) at 2 weeks, and approximately 84 % of H3K27ac-marked SEs overlapped with hypomethylated DMRs (Fig. 2a, b). Gene ontology analysis demonstrated that genes associated with these hypomethylated SEs were significantly enriched for tissue regenerative and transcriptional programs, including epithelial development, wound-healing, cell–cell adhesion, and DNA-binding transcriptional processes (Fig. 2c and Supplementary Fig. 2c,). Collectively, these findings indicate that, in this YAP1-driven epigenetic carcinogenesis model, enhancer DNA hypomethylation represents a key mechanism underlying the irreversible transition from dysplasia to carcinoma *in situ*.

**Figure 2.**
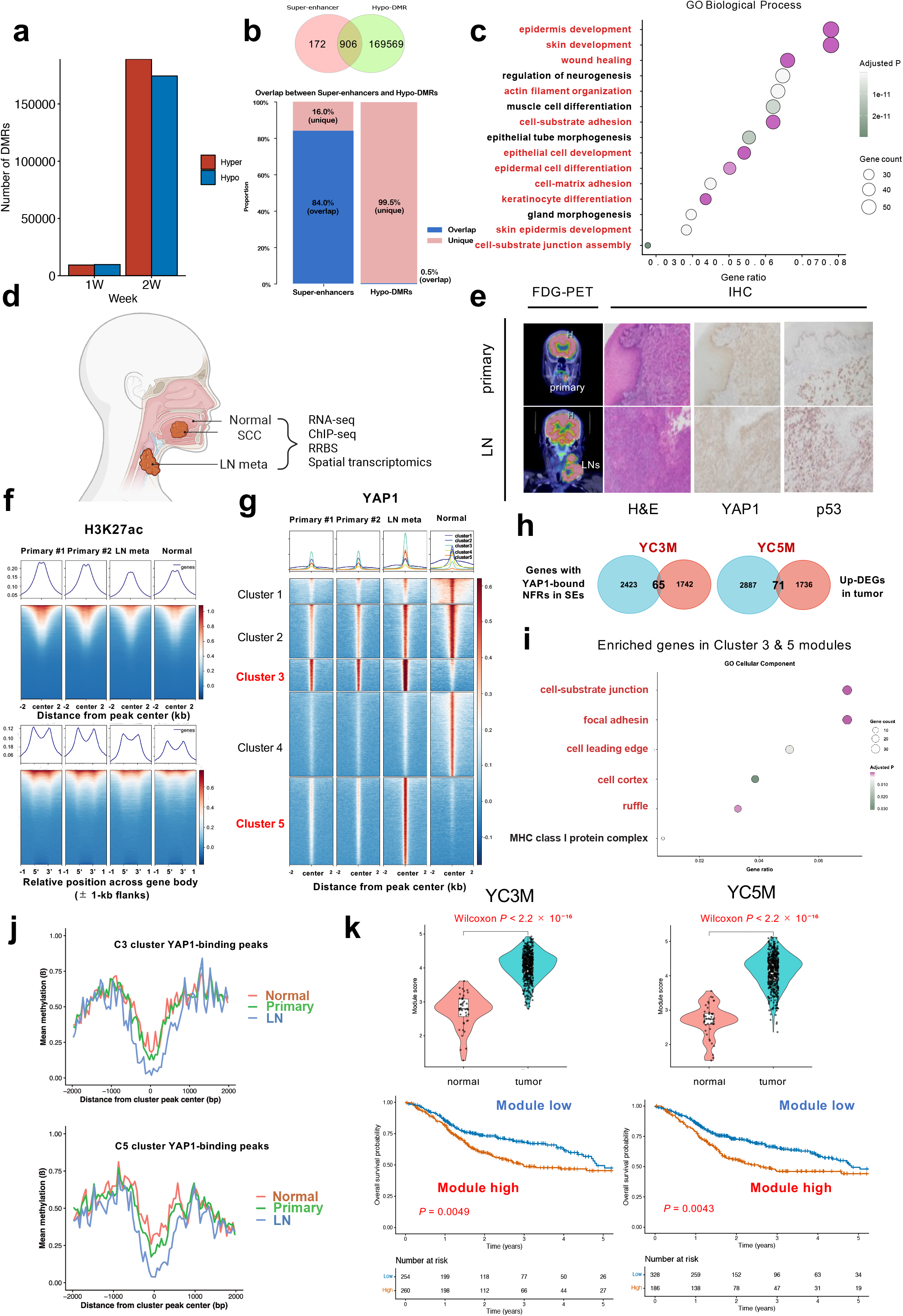
Context-dependent redistribution of YAP1 and enhancer reprogramming during human HNSCC evolution. **a** Numbers of differentially methylated regions (DMRs) identified by WGBS in *iMob1*DKO cells at 1 and 2 weeks after tamoxifen treatment. Hypomethylated DMRs markedly increased at 2 weeks. **b** Venn diagram and proportion plot showing overlap between H3K27ac-marked super-enhancers (SEs) and hypomethylated DMRs at 2 weeks. **c** Gene Ontology (GO) Biological Process enrichment of genes associated with hypomethylated SEs. **d** Multiregional sampling of human HNSCC specimens comprising normal epithelium, primary tumors, and lymph node (LN) metastases for RNA-seq, ChIP-seq, reduced representation bisulfite sequencing (RRBS), and spatial transcriptomics. Created in BioRender. https://BioRender.com/pm4mmzq. **e** FDG-PET images showing faster growth of LN metastases than the primary tumor; corresponding hematoxylin and eosin and YAP1/p53 immunohistochemical staining. **f** H3K27ac ChIP-seq average signal profiles and heatmaps around peak centers (±2 kb) and across gene bodies with 1-kb flanking regions. **g** YAP1 ChIP-seq heatmap centered on binding peaks (±2 kb) in multiregional samples. **h** Integration of YAP1-associated SE profiles, nucleosome-free regions, and RNA-seq identifies overexpressed gene modules associated with YAP1-bound SEs in clusters 3 (YC3M, 65 genes) and 5 (YC5M, 71 genes). **i** GO Biological Process enrichment of YC3M and YC5M genes, highlighting cell–matrix interactions. In **c** and **i**, the x-axis indicates gene ratio, circle size indicates gene count, and color indicates Benjamini–Hochberg-adjusted *P* values. **j** Average RRBS DNA methylation profiles around cluster 3 and cluster 5 YAP1-binding peak centers (±2 kb), showing progressive hypomethylation from normal mucosa through primary tumors to matched LN metastases. **k** Violin plots showing increased YC3M and YC5M activities in TCGA HNSCC tumors versus normal tissues (Wilcoxon rank-sum test, both *P* < 2.2 × 10□¹□). Kaplan–Meier analyses show poorer overall survival with high YC3M or YC5M activity (log-rank test, *P* = 0.0049 and 0.0043, respectively). See Source Data for **j** and **k**.

### Context-dependent redistribution of YAP1 and enhancer rewiring during human HNSCC evolution

While the mouse model suggests that YAP1 activation alone can induce epigenetic carcinogenesis, human HNSCC develops within a genetic context, frequently involving *TP53* mutations ^4^. To bridge this gap and assess the interplay between genetic alterations and YAP1-driven epigenetic remodeling, we conducted multi-omics analyses of multiregional (normal tissue, primary tumors, and LN metastases) human HNSCC samples exhibiting *TP53* mutations (loss-of function Y234C) and high YAP1 activity (Fig. 2d, e). Clinically, these tumors showed an exceptionally aggressive metastatic phenotype, characterized by more rapid growth in LNs than in the primary tumor as demonstrated by FDG-PET imaging (Fig. 2e) and early systemic dissemination to distant organs shortly after initial surgery. We performed RNA sequencing (RNA-seq), ChIP-seq for the constitutive heterochromatin mark H3K9me3, the active enhancer mark H3K27ac, and YAP1, as well as reduced representation bisulfite sequencing (RRBS). In contrast to previous findings in pancreatic cancer ^45^, metagene analysis of H3K9me3 ChIP–seq signals across gene bodies and flanking regions revealed relative stability of constitutive heterochromatin domains during HNSCC evolution (Supplementary Fig. 2d). By contrast, the peak intensities of H3K27ac and YAP1 exhibited marked differences among normal tissue, primary tumors, and LN metastases. Accordingly, YAP1 cluster 3 was prominent in both primary tumors and LNs, whereas YAP1 cluster 5 was uniquely observed in LN metastases (Fig. 2f, g). After confirming significantly elevated expression of cluster 3 (n = 1,674) and 5 (n = 1,818) genes in tumors and LN metastases compared with normal tissue (*P* < 0.001; supplementary Fig2e), we integrated these results with SE profiling, a nucleosome-free regions (NFRs) estimation algorithm, and RNA-seq data (Fig. 2h). Consequently, we identified 65 and 71 YAP1-associated SE loci in cluster 3 and 5, respectively, and thereby defined their upregulated target genes, which we termed YAP1 associated cluster 3 module (YC3M) and cluster 5 module (YC5M) (Fig.2h, and Supplementary Data 2). Functional enrichment analysis revealed that these module genes were significantly associated with cell–matrix interactions (Fig. 2i), consistent with the enhanced metastatic capacity to LNs. Notably, RRBS analysis demonstrated progressive DNA hypomethylation surrounding C3 and C5 YAP1-binding peaks from normal mucosa to primary tumors and further to LN metastases (Fig. 2j). Moreover, super-enhancers harboring C3 or C5 YAP1-binding peaks were significantly hypomethylated relative to background super-enhancers in all three tissue types (Supplementary Fig. 2f; P < 0.0001), consistent with findings in *iMob1*DKO cells (Fig.2a). Analysis of TCGA datasets further demonstrated that the activities of both modules were significantly higher in tumor samples than in normal tissues (*P* < 2.2 × 10^-16^), and that patients with high module activity exhibited significantly poorer overall survival (*P* = 0.0049 and *P* = 0.0043 for YC3M and YC5M, respectively; Fig. 2k). Collectively, these results indicate that YAP1 dynamically redistributes to distinct enhancer landscapes in a context-dependent manner across the normal–carcinogenesis–metastasis continuum, thereby promoting malignant transcriptional reprogramming through SE formation and enhancer DNA hypomethylation.

### YAP1 orchestrates super-enhancer formation and establishes a feedforward oncogenic circuit in HNSCC

To mechanistically interrogate how YAP1 reshapes enhancer architecture and modulates target gene expression, we performed H3K27ac and YAP1 ChIP–seq in combination with RNA-seq using a doxycycline-inducible YAP1 overexpression system in YAP1-low SCC9 cells and a YAP1 knockdown (shYAP1) system in YAP1-high HSC4 cells, both of which we established previously^19^. In HSC4 cells, genome-wide occupancy analysis revealed that approximately 90% of YAP1-binding sites were located outside promoter regions, predominantly within distal intergenic and intronic regions (Supplementary Fig. 3a, left). Consistent with this distribution, YAP1 binding closely overlapped H3K27ac-marked regions, indicating preferential binding to active distal enhancers (Supplementary Fig. 3a, right). Functional annotation of YAP1-bound enhancers showed significant enrichment for wound healing, epithelial cell development, and cell–matrix interaction programs (Supplementary Fig. 3b), closely mirroring the functional enrichment observed for hypomethylated DMR-associated genes (Fig. 2c). Integration of ChIP-seq data with RNA-seq data identified 19 direct YAP1 target genes—designated the YAP1 progression module (YPM)—including *RIMS3, DHCR24, AK4, DDAH1, PTPN14, ADK, CD59, DENND5B, KRT80, CKAP2, MBNL2, COL4A2, BUB1B, TTBK2, USO40, PHF20, IL6, MTURN,* and *CUX1* (Fig. 3a). Clinically, elevated YPM activity was significantly associated with poorer overall survival in TCGA HNSCC cohorts (*P* = 0.031; Fig. 3b), supporting its functional relevance in tumor progression. Using the ROSE algorithm, we identified 342 super-enhancer (SE)-associated genes in HSC4 cells, among which YPM genes were prominently represented (Fig. 3c). Notably, YAP1 occupancy was markedly enriched at SEs compared to typical enhancers (334/342 vs 3047/6049; Fisher’s exact test, odds ratio = 44.1, *P* < 2.2 × 10□^16^; Fig.3d), demonstrating a strong bias toward SE architecture. Consistently, YAP1 silencing reduced global H3K27ac signal intensity and led to a ∼43% decrease in SE number (Supplementary Fig. 3c), indicating that YAP1 is required for both the establishment and maintenance of SEs. Genes linked to YAP1-dependent SEs were enriched for cytoskeletal organization, epithelial differentiation, and cell–matrix interaction pathways (Fig. 3f), in agreement with our previous observations (Fig. 1b, 2c, and Supplementary Fig 3b). Among the YAP1-associated SE target genes, IL6 attracted particular attention. IL6 is a major HNSCC-causing cytokine abundant in the HNSCC TME ^46^ and its prognostic relevance was confirmed in TCGA dataset (*P* = 0.0075; Fig. 3f). In addition, IL6 has been reported to activate YAP1 through Src-mediated phosphorylation at Y357 ^36^. In YAP1-high HSC4 cells, IL6 was identified as a target of YAP1-associated SE (Fig. 3g), and siRNA-mediated suppression of IL6 significantly reduced YAP1 Y357 phosphorylation and inhibited cell proliferation (Fig. 3h). Conversely, in YAP1-low Cal27 cells, exogenous IL6 increased YAP1 Y357 phosphorylation and promoted cell growth (Supplementary Fig. 3d). These findings support the presence of a super-enhancer–driven YAP1–IL6 feedforward loop that contributes to the initiation and progression of HNSCC in a subset of tumors (Fig. 3i).

**Figure 3.**
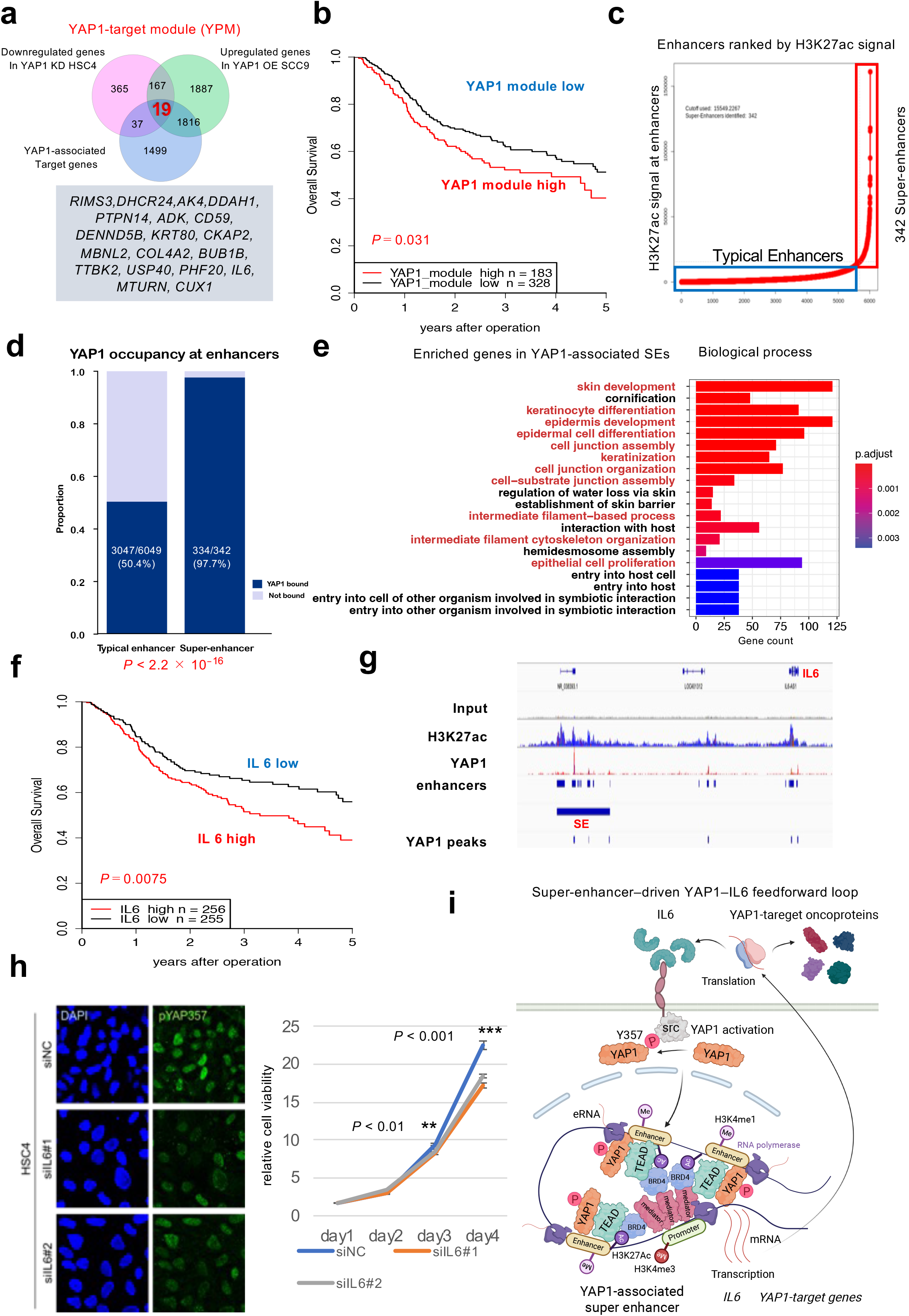
YAP1 orchestrates super-enhancer formation and establishes a feedforward oncogenic circuit in HNSCC. **a** Integration of H3K27ac and YAP1 ChIP–seq and RNA-seq analyses identified 19 direct YAP1 target genes, defined as the YAP1 progression module (YPM). Venn diagram showing overlap between YAP1-bound enhancer-associated genes and differentially expressed genes. **b** Kaplan–Meier analysis of overall survival in the TCGA HNSCC cohort stratified by YPM activity (ssGSEA score). Elevated YPM activity was associated with poorer prognosis (*P* = 0.031, log-rank test). **c** Super-enhancer (SE) landscape defined by H3K27ac ChIP–seq using the ROSE algorithm. **d** Proportion plot comparing YAP1 occupancy between typical enhancers and SEs. **e** Functional annotation (GO Biological Process) of genes reduced genes targeted by YAP1-associated SEs. Bars indicate the number of differentially expressed genes assigned to each GO Biological Process term. Colors represent Benjamini–Hochberg-adjusted *P* values. **f** Kaplan–Meier analysis of overall survival in the TCGA HNSCC cohort stratified by the levels of *IL6* mRNA expression (*P* = 0.0075, log-rank test). **g** Genome browser tracks at the IL6 locus demonstrating YAP1 binding and SE-associated H3K27ac enrichment in HSC4 cells. **h** Functional validation of the YAP1–IL6 feedforward loop. IL6 knockdown reduced YAP1 Y357 phosphorylation in YAP1-high HSC4 cells (left) and significantly suppressed cell proliferation compared with control siRNA (right). Data are presented as mean ± SD. Comparisons between control siRNA and each IL6-targeting siRNA were performed using two-sided Student’s *t*-tests at each time point. *P* < 0.01 (day 3) and *P* < 0.001 (day 4). **i** Schematic model illustrating a super-enhancer–driven YAP1–IL6 feedforward loop that amplifies oncogenic transcriptional programs during HNSCC progression. Created in BioRender. https://BioRender.com/w77armt. See Source Data for i.

### Stage-specific recruitment of partner transcription factors to YAP1-associated super-enhancers

To explain the mechanism underlying the context-dependent redistribution of YAP1 to SEs of target genes (Fig. 2h), we hypothesized that, in addition to the canonical partners TEAD and AP-1, YAP1 may cooperate with non-canonical transcription factors during HNSCC evolution. We first searched for candidate partner TFs relevant to the carcinogenesis stage, where master regulators of the squamous epithelial lineage are expected to play key roles in promoting self-renewal and tissue-regenerative transcriptional programs. Based on literature curation, we identified two candidate master transcription factors—FOXM1 and ΔNp63 (p40)—both of which have been reported to directly interact with YAP1 and promote tumorigenesis.^47,48^ ^49,50^ We then examined gene sets using publicly available FOXM1- and ΔNp63-specific human ChIP–seq datasets ^51,52^ and found that, among the YAP1 target genes within the YCM module (Fig. 1C), *TK1, TOP2A, CENPF, CDC20,* and *HMMR* were FOXM1 targets, whereas *TK1, FSCN, AXL, and HMMR* were ΔNp63 targets. Using 17 human HNSCC specimens, we confirmed that YAP1, FOXM1, and ΔNp63 proteins exhibited similar spatial distribution patterns, expanding from the basal/stem–progenitor compartment in normal epithelium to widespread expression across tumor cells (Fig. 4a). Triple-positive tumors (YAP1⁺/FOXM1⁺/ΔNp63⁺) were significantly enriched in the YAP1-positive group (6/7 vs 0/10; Fisher’s exact test, *P* = 5.66 × 10⁻^4^; Haldane-corrected OR = 91.0, 95% CI 3.20–2585.49; Fig.4a). Consistent with these observations, a recent HNSCC study of our group has demonstrated interaction between YAP1 and FOXM1^22^, and the essential role of ΔNp63 in squamous carcinogenesis is well established ^48^. Collectively, these findings suggest that, during the early carcinogenic process, YAP1 cooperates with lineage-defining master transcription factors—FOXM1 and ΔNp63—to activate an intrinsic tissue-regenerative transcriptional program.

**Figure 4.**
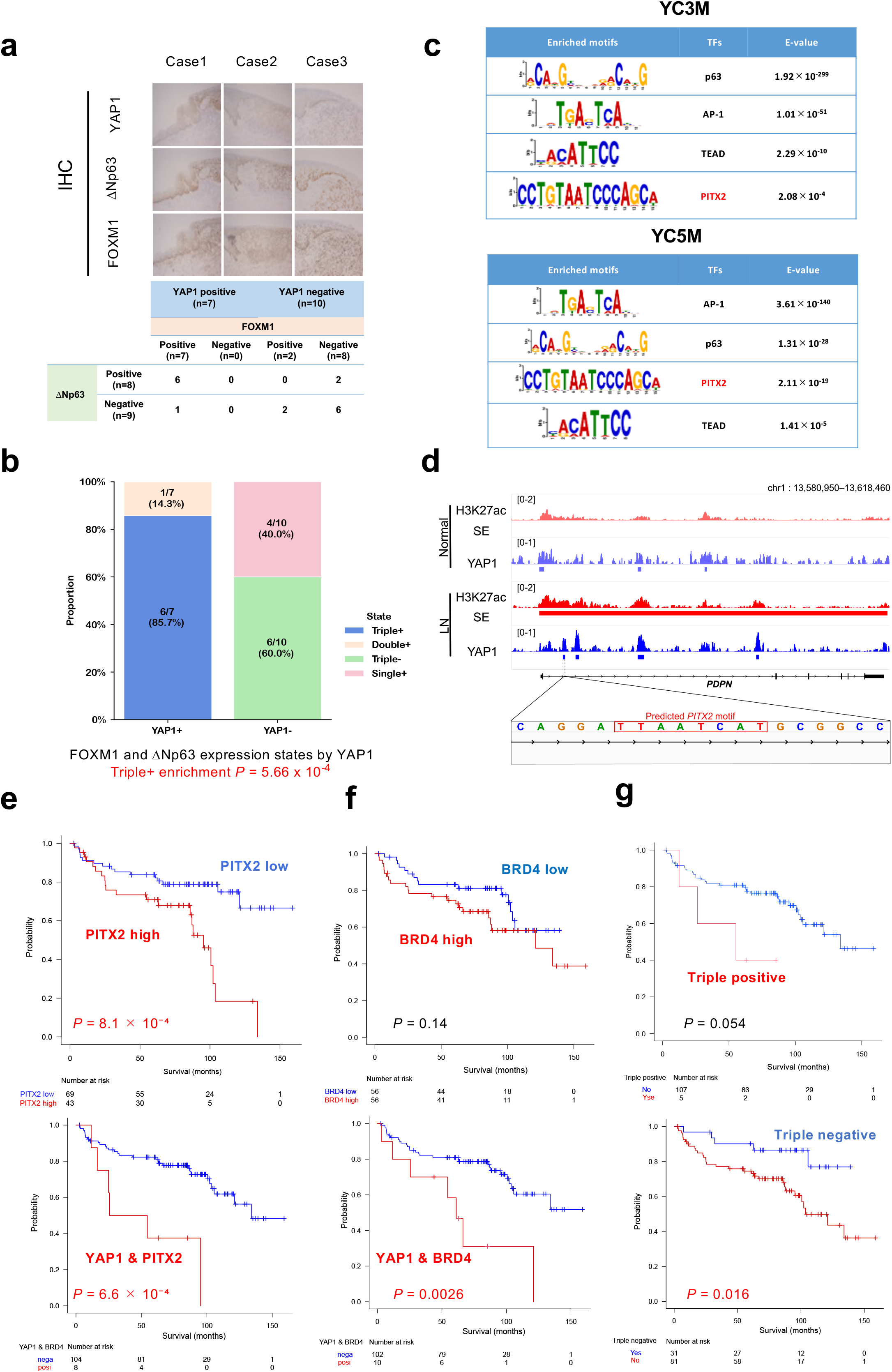
Stage-specific transcription factor partnerships at YAP1-associated enhancers during HNSCC progression. **a** Representative immunohistochemical analysis of YAP1, FOXM1, and ΔNp63 expression in HNSCC specimens (n = 17). The table summarizes the frequencies of FOXM1- and ΔNp63-positive cases according to YAP1 expression status. **b** Association of YAP1 expression with FOXM1 and ΔNp63 expression patterns in HNSCC specimens. Stacked bar chart showing the proportions of triple-positive (YAP1 /FOXM1□/ΔNp63□), double-positive, and single-positive tumors according to YAP1 expression status (n = 17). Triple-positive tumors were significantly more frequent in the YAP1-positive group than in the YAP1-negative group (Fisher’s exact test, *P* = 5.66 x 10^-4^). **c** Selected transcription factor motifs in YAP1-bound super-enhancers associated with YC3M and YC5M. The p63 and AP-1 motifs met the significance threshold (*E* < 1 × 10□□) in both modules. The PITX2 motif met this threshold only in YC5M, and the TEAD motif only in YC3M.. **d.** Genome browser tracks at the *PDPN* locus demonstrating YAP1 binding and SE-associated H3K27ac enrichment in human LN samples with PITX2 binding sequence. **e, f** Kaplan–Meier analyses of overall survival in the HNSCC cohort (n = 112). PITX2 protein expression alone or co-expression with YAP1 was associated with worse prognosis (*P* = 8.1 × 10□□ and *P* = 6.6 × 10□□, respectively; log-rank test). BRD4 expression alone was not significantly associated with prognosis (*P* = 0.14), whereas co-expression of BRD4 with YAP1 was associated with worse prognosis (*P* = 0.0026; log-rank test). **g** Kaplan–Meier analyses according to combined YAP1, PITX2, and BRD4 expression. Triple-positive tended to have shorter overall survival (*P* = 0.054), whereas triple-negative tumors showed significantly better overall survival (*P* = 0.016; log-rank test). See Source Data for **e, f**, and **g**.

We next investigated candidate YAP1 partner TFs in the progression setting. Motif discovery analyses were performed to identify DNA-binding motifs associated with YAP1-bound SEs, using datasets derived from YAP1 target genes including YPM in human HNSCC cell lines (Fig. 3a) as well as YC3M and YC5M genes identified in human samples (Fig. 2h). Motif analysis identified p63 and PITX2 (paired-like homeodomain 2) as candidate YAP1 partners alongside AP-1 and TEAD (Fig. 4c and Supplementary Fig. 4a). The AP-1 and p63 motifs were significantly enriched in both YC3M and YC5M. Notably, the TEAD motif met the significance threshold only in YC3M (*E* = 2.29 × 10□¹□), whereas the PITX2 motif met this threshold only in YC5M (*E* = 2.11 × 10□¹□) (Fig. 4c). An independent analysis also identified the PITX2 motif among the top enriched motifs associated with genes upregulated in lymph node metastases relative to paired primary tumors (RNA-seq, n = 10) (Supplementary Fig. 4b). PITX2 has been reported as an unfavorable prognostic factor in esophageal cancer, where it promotes stemness, and as a regenerative transcription factor that can directly interact with YAP1 ^53,54^. Consistently, immunoprecipitation assays confirmed that PITX2 physically associates with YAP1 in HNSCC cell lines (Supplementary Fig. 4c), as confirmed in our recent study ^22^. At the enhancer locus of *PDPN*, a representative YC5M gene, a YAP1-associated super-enhancer containing a PITX2-binding motif was identified in human lymph node metastasis but was absent in matched normal tissue (Fig.4d). In TCGA datasets, higher PITX2 mRNA expression was associated with poorer survival in patients with HNSCC (Supplementary Fig. 4d), and PITX2 protein expression—either alone (*P* = 8.1 × 10□□) or co-expressed with YAP1(*P* = 6.6 × 10□□)—predicted worse clinical outcomes in our cohort (n = 112) (Fig. 4e and Supplementary Fig. 4e). Notably, the prognostic impact of PITX2 expression was greater than that of BRD4 (*P* = 0.14 or 0.0026), a core component of SEs across multiple cancers including HNSCC (Fig. 4f and Supplementary Fig. 4e) ^25,34^. Patients with triple-positive tumors tended to have shorter overall survival than the other patients (*P* = 0.054), whereas those with triple-negative tumors had significantly longer overall survival than the other patients (*P* = 0.016) (Fig. 4g). In addition, we previously demonstrated that YAP1 promotes the acquisition of stemness in HNSCC through cooperation with SOX2 ^20^. Collectively, our data support a model in which YAP1-associated SEs act as dynamic regulatory hubs that recruit stage-specific client transcription factors, thereby orchestrating context-dependent enhancer reprogramming during HNSCC progression.

### The YAP1–PITX2 axis drives partial EMT and CAF reprogramming during lymph node metastasis

Given the potential role of the YAP1–PITX2 axis in HNSCC progression, particularly in LN metastasis, we further investigated their functional relevance using recently published single-cell RNA-seq datasets of HNSCC^55^. This landmark study demonstrated that LN metastasis is driven by interactions between partial EMT (p-EMT) cells and CAFs reprogrammed by TGF-β signaling, and identified a distinctive 15-gene p-EMT signature (Supplementary Data 3). Consistent with this framework, downregulation of PITX2 in HSC4 cells significantly reduced the expression of five p-EMT signature genes (*TGFb1, LAMC2, ITGA5, PDPN* and *CDH13*), as well as stemness-associated markers (*CD44, CD24* and *ALDH1A3*) and hypoxia related gene *HIF1A* (Fig. 5a). Interestingly, *TGFb1, LAMC2, PDPN,* and *HIF1A* were included in the LN-specific YC5M genes (Supplementary Data 2). Gene ontology analysis of downregulated DEGs revealed enrichment for pathways related cellular motility and adhesion (Supplementary Fig. 5a). We previously demonstrated that PITX2 silencing impaired cell proliferation, sphere-forming capacity, and migration in HNSCC cells ^22^. Consistent with these findings, independent sphere-formation assays performed in the present study showed that PITX2 silencing significantly reduced sphere-forming capacity (Supplementary Fig. 5b), further supporting a functional role for PITX2 in maintaining the sphere-forming capacity of YAP1-active HNSCC cells. Uniform Manifold Approximation and Projection (UMAP) embedding of the single-cell RNA-seq data ^55^ revealed that HNSCC cells with high PITX2 expression tended to exhibit elevated YPM activity (Fig. 5b). We next performed Expression Enrichment Matrix (EEM) analysis ^56^ using TCGA datasets to identify synchronized transcriptional programs associated with the YAP1 enhancer target genes identified in Fig.3 (Fig. 5c). The analysis revealed coordinated programs encompassing cell cycle, EMT, matrix remodeling, YAP1-associated programs, TGF-β signaling, immune response, and inflammation. Notably, YAP1 enhancer target activity was associated with EMT and matrix-remodeling programs, together with IL6 and TGF-β signaling, particularly in tumors with relatively low cell-cycle activity, consistent with an invasive and stromal-remodeling transcriptional state. Further supporting this observation, YPM activity in TCGA HNSCC samples was positively correlated with both the p-EMT signature (*R* = 0.53, *P* < 2.2 × 10□¹□, Fig. 5d) and a TGF-β–induced CAF signature defined in Supplementary Data 3 (*R* = 0.42, *P* < 2.2 × 10□¹□, Fig. 5e)^57,58^. Collectively, these findings establish the YAP1–PITX2 complex as a central reprogramming driver of p-EMT tumor cells and CAFs, thereby promoting LN metastasis in HNSCC.

**Figure 5.**
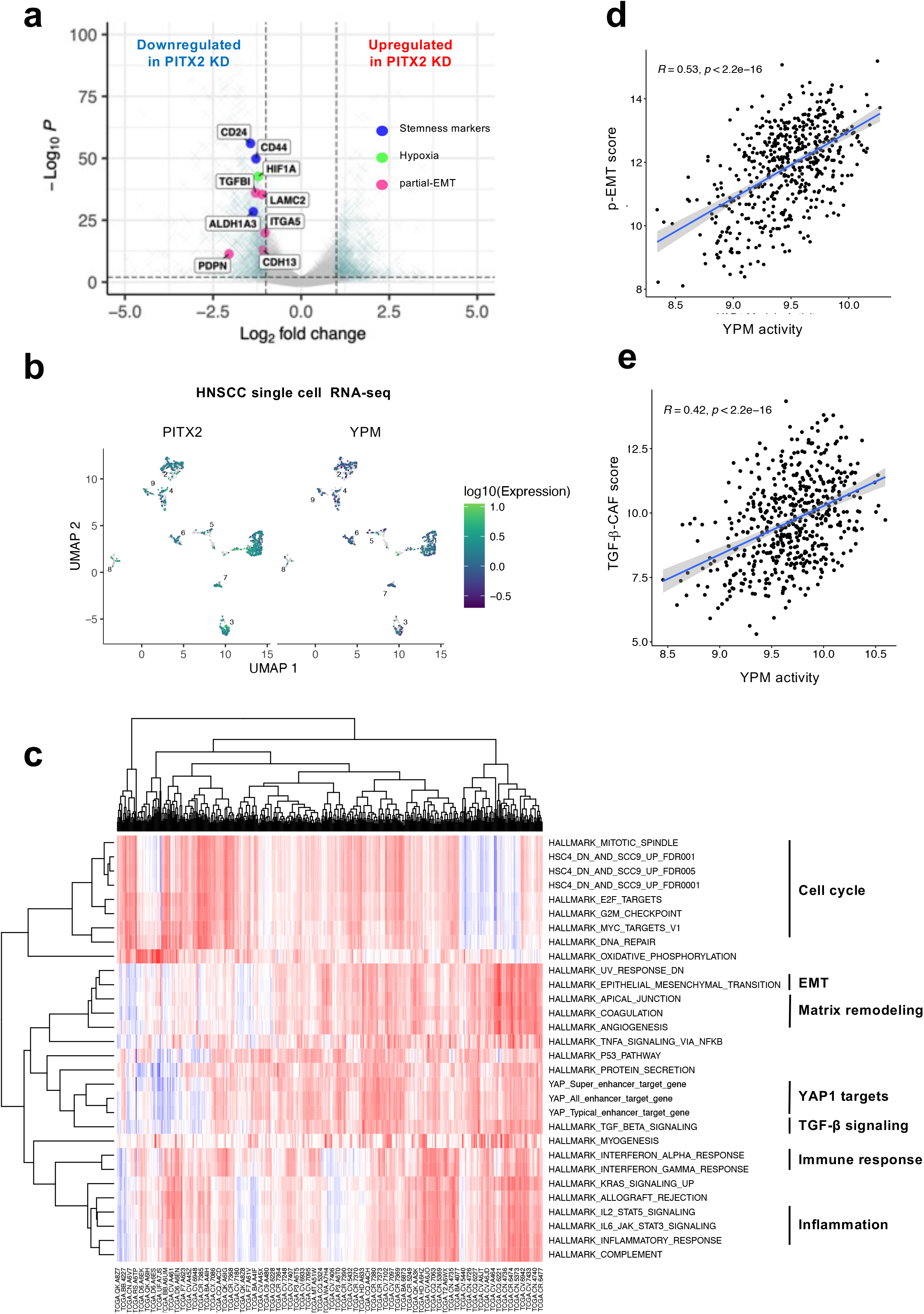
The YAP1–PITX2 axis promotes partial EMT and CAF-associated transcriptional programs during lymph node metastasis. **a** Volcano plot showing differentially expressed genes following PITX2 knockdown in HSC4 cells. p-EMT signature genes (TGFB1, LAMC2, ITGA5, PDPN, CDH13), stemness markers (CD44, CD24, ALDH1A3), and HIF1A are highlighted. **b** UMAP embedding of published HNSCC single-cell RNA-seq data□□. Cells are colored according to PITX2 expression (left) or YAP1 progression module (YPM) activity (right), demonstrating co-enrichment. **c** Expression Enrichment Matrix (EEM) analysis of TCGA HNSCC samples showing coordinated transcriptional programs associated with YAP1 enhancer target genes identified in Fig. 3. Gene sets were grouped into functional categories including cell cycle, inflammation, EMT, matrix remodeling, YAP1 targets, TGF-β signaling, and immune response. **d** Correlation between YPM activity and the p-EMT signature in TCGA HNSCC cohort (R = 0.53, P < 2.2 × 10□^16^). **e** Correlation between YPM activity and a TGF-β–induced CAF signature in TCGA dataset (R = 0.42, P < 2.2 × 10□^16^).

### Spatial transcriptomics identifies a YAP1/PITX2-associated metastatic niche characterized by p-EMT, CAF interactions, and immune suppression

In our final set of experiments, we sought to further investigate the role of the YAP1/PITX2 axis in lymph node metastasis and TME remodeling, given the potent immunosuppressive and pro-fibrotic functions of IL6 and TGF-β signaling in HNSCC and other cancers ^40,58^. To this end, we performed Visium spatial transcriptomic analysis using the same FFPE multiregional HNSCC specimens that were subjected to the integrated multi-omics analyses described above where YAP1/PITX2 axis plays an essential role in tumor evolution (Figs. 2d, and 4c, d). Because YAP1-driven transcriptional reprogramming was closely associated with nuclear YAP1 protein expression (Fig. 1), we selected YAP1-high regions of interest (ROIs) from primary tumors and matched lymph node metastases for spatial transcriptomic profiling. For comparison, we selected areas of normal epithelium within the primary tumor ROI labeled T2 in Fig. 6a. To validate the appropriateness of ROI selection, we first calculated YAP1 target gene scores—YCM, YPM, YC3M, and YC5M—across the analyzed regions. YC3M, YC5M, and YPM activities increased progressively from normal epithelium to primary tumors and then to lymph node metastases (*P* < 2.2 × 10□^¹□^ for each pairwise comparison). YCM activity differed only modestly between normal epithelium and primary tumors (*P* = 0.038), and was highest in lymph node metastases (Supplementary Fig. 6a). Constitutive expression of YCM genes in normal basal stem cells may explain the modest difference in YCM activity between normal epithelium and primary tumors. These findings confirmed that the selected ROIs captured spatial regions with active YAP1-driven transcriptional programs.

**Figure 6.**
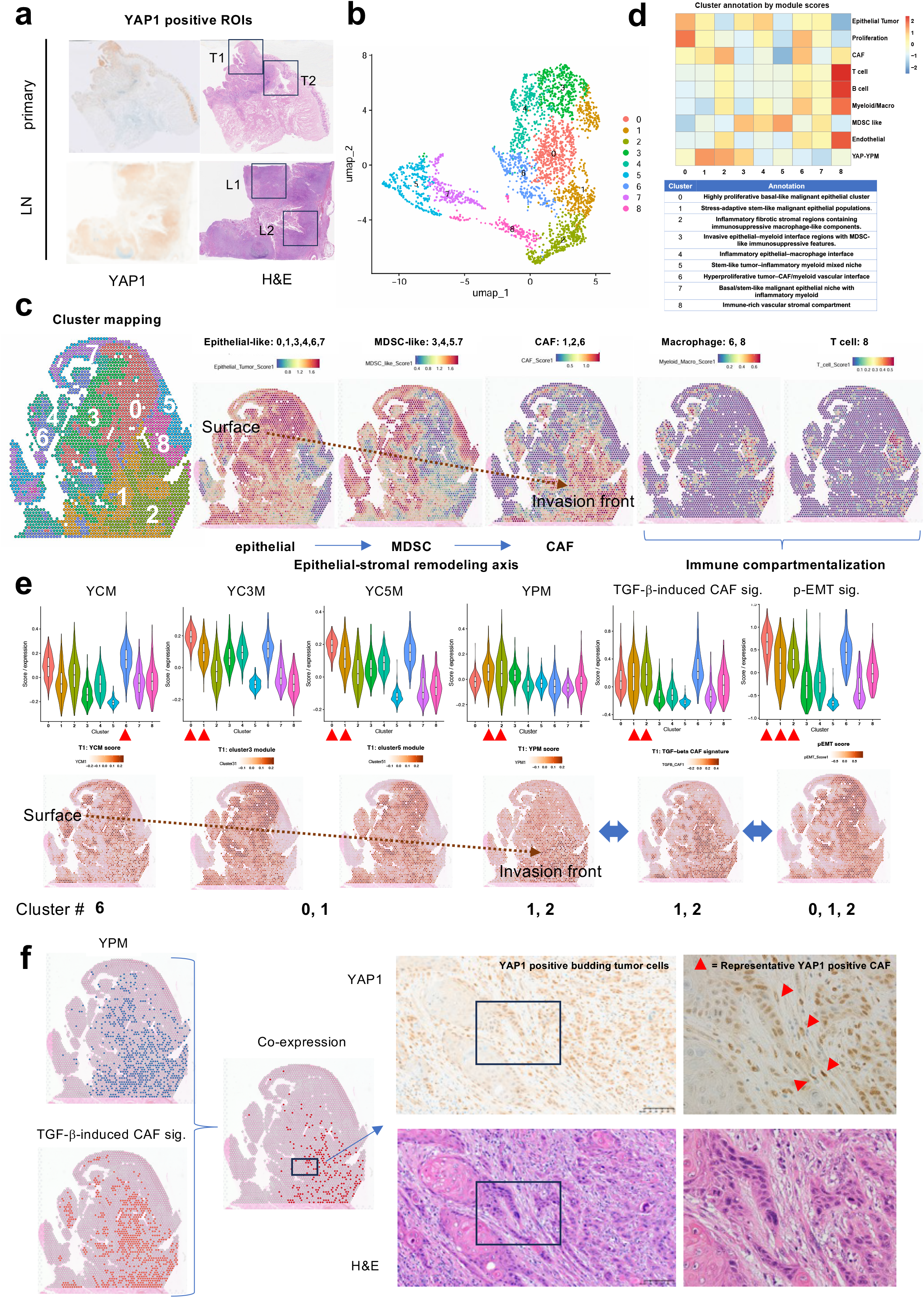
Spatial transcriptomic analysis reveals YAP1-associated epithelial, stromal, and immune ecosystem remodeling during HNSCC progression. **a** Selection of regions of interest (ROIs) for Visium spatial transcriptomic profiling of HNSCC specimens previously characterized by multi-omics analysis. YAP1-high ROIs were selected from primary tumors and matched lymph node metastases. Areas of normal epithelium were selected from within the primary tumor ROI labeled T2. **b** UMAP visualization of Visium spots identifying nine spatial ecosystems (clusters 0–8). **c** Spatial distribution of the nine transcriptomic clusters and major cellular ecosystems across representative HNSCC tissue sections. Clusters were annotated based on canonical marker genes, spatial localization, and cell2location-inferred cellular composition. Representative tissue sections reveal a spatial epithelial–stromal remodeling axis extending from the tumor surface to the invasive front, characterized by a progressive transition from epithelial to MDSC-like and CAF-rich ecosystems. Macrophage-rich and T-cell-rich ecosystems formed distinct spatial compartments outside the remodeling axis. **d** Heat map of nine clusters according to the estimated cellular lineages (upper). Annotation of the nine spatial ecosystems based on lineage markers, inferred cellular composition, and spatial organization (lower). **e** Spatial epithelial–stromal remodeling axis and YAP1-associated transcriptional evolution during tumor invasion. Clusters 6, 0, 1, and 2 formed a continuous spatial axis extending from superficial tumor regions toward the invasive front. Along this remodeling axis, YAP1-associated transcriptional programs progressively evolved from YCM to YC3M, YC5M, and ultimately YPM. TGF-β-induced CAF and p-EMT signatures were selectively enriched at the invasive front. **f** Histopathological validation of the fibrotic metastatic ecosystem. Representative spatial maps showing YPM activity, TGF-β-induced CAF signature, and their co-localization (left). The corresponding histological sections demonstrate YAP1-positive budding tumor cells invading the surrounding stroma in close association with YAP1-positive cancer-associated fibroblasts (CAFs) within regions exhibiting concurrent high YPM activity and TGF-β-induced CAF signatures (right). See Source Data for e.

UMAP clustering of Visium spots identified nine transcriptionally distinct spatial ecosystems (clusters 0–8) (Fig. 6b). Cluster identities were assigned using canonical lineage markers (Supplementary Data 3) together with spatial distributions of inferred cellular lineages (Supplementary Fig. 6b–d and Fig. 6c, d). Rather than representing individual cell types, these clusters corresponded to composite tumor–stromal niches defined by characteristic lineage compositions and biological programs, including proliferative malignant epithelial regions, invasive epithelial–myeloid interfaces, fibrotic stromal compartments, and immune-rich vascular niches (Fig.6d). These analyses revealed extensive ecosystem-level heterogeneity within YAP1-active tumors, suggesting coordinated remodeling of epithelial, stromal, and immune compartments during tumor progression.

Among the identified ecosystems, we focused on clusters 6, 3, 0, 1, and 2, which formed a spatial continuum extending from superficial tumor regions toward the invasive front (Fig. 6c). Along this trajectory, the cellular composition progressively shifted from epithelial-rich tumor ecosystems to MDSC-associated intermediate ecosystems and ultimately to CAF-enriched invasive ecosystems, revealing a spatial epithelial–stromal remodeling axis during tumor invasion (Fig. 6c). These invasive ecosystems remained largely depleted of lymphoid populations, suggesting spatial compartmentalization of immune responses during YAP1-driven tumor evolution (Fig. 6c). Along this ecosystem gradient, YAP1-associated transcriptional programs evolved sequentially from YCM to YC3M and YC5M, ultimately converging on YPM, consistent with stepwise transcriptional reprogramming during tumor progression and invasion (Fig. 6e). Notably, YPM-high regions were consistently located adjacent to, and partially overlapped with, areas enriched for a TGF-β-induced CAF signature at the invasive front, particularly within clusters 1 and 2 (Fig. 6e). This spatial arrangement suggested the presence of a specialized invasive niche in which YAP1-driven tumor cells and activated fibroblasts engage in reciprocal interactions. Consistent with this interpretation, p-EMT scores were significantly elevated in epithelial clusters 0, 1, and 2, corresponding to regions undergoing active invasion (Fig. 6e). Given our identification of PITX2 as a progression-associated YAP1 partner transcription factor in identical samples (Fig.4c), these findings support a model in which YAP1/PITX2-mediated transcriptional reprogramming promotes a p-EMT state that cooperates with TGF-β-activated CAFs to facilitate metastatic progression. Supporting this notion, histopathological examination demonstrated that regions exhibiting concurrent YPM and TGF-β-induced CAF signatures were characterized by budding YAP1-positive tumor cells invading into the surrounding stroma, which were closely encircled by YAP1-positive CAFs (Fig. 6f). Together, these findings identify a YAP1/PITX2-associated fibrotic metastatic ecosystem in which p-EMT tumor cells and activated CAFs cooperatively establish an invasive niche that supports metastatic dissemination.

We next investigated how immune ecosystems were spatially organized relative to invasive YAP1-active niches. Clusters 3, 4, 5, and 7 represented epithelial–myeloid ecosystems characterized by expression of S100A8, S100A9, ARG1, IL1B, CXCL8, and LILRB2 (Supplementary Data 3), consistent with MDSC-like immunosuppressive programs (Fig. 6b-d and supplementary Fig. 6d). Notably, these ecosystems were spatially segregated from YPM-high invasive fronts and instead accumulated within and around superficial tumor islands (Fig. 6c, e). In contrast, cluster 8 constituted an immune-rich vascular stromal ecosystem (Fig.6d) characterized by high IL6 expression (Fig.7a) and enrichment signatures (Supplementary Data 3) of leukocyte (Fig.7b), endothelial, B-cell (supplementary Fig.6d), and CD8⁺ T-cell markers (Fig.7c). Notably, CD8⁺ T cells were largely confined to this ecosystem and displayed an exhaustion-like transcriptional phenotype (Supplementary Data 3 and Fig.7c). These observations suggest that YAP1-active tumors establish a spatially compartmentalized immunosuppressive ecosystem, in which MDSC-rich tumor niches, fibrotic invasive fronts, and IL6-associated exhausted T-cell compartments occupy distinct but interconnected spatial territories.

Collectively, spatial transcriptomic analyses revealed that YAP1-driven tumor evolution is accompanied by the emergence of functionally distinct metastatic ecosystems. These included a YAP1/PITX2-associated p-EMT and CAF-rich invasive niche, MDSC-dominated epithelial–myeloid compartments, and an adjacent IL6-rich immune-restricted vascular ecosystem harboring exhausted CD8⁺ T cells. Together, these findings indicate that enhancer rewiring downstream of YAP1 extends beyond malignant epithelial cells and is associated with coordinated stromal and immune remodeling that culminates in the establishment of a fibrotic and immunosuppressive metastatic microenvironment.

Although these observations were derived from a representative multiregional HNSCC specimen, they suggested coordinated epithelial, stromal, and immune remodeling downstream of YAP1 activation. To determine whether these findings were generalizable, we next examined the relationship between YAP1-associated transcriptional programs and tumor immune phenotypes in the TCGA HNSCC cohort using a public molecular functional tumor portrait platform (https://science.bostongene.com/tumor-portrait/)^59^. Fibrotic tumors exhibited significantly higher YPM and YCM activities than desert tumors, while immune-enriched fibrotic tumors showed significantly higher YPM and YCM activities than immune-enriched non-fibrotic tumors (Fig. 7d). Together, these findings indicate that YAP1-driven transcriptional reprogramming is preferentially associated with fibrotic, immunosuppressive, and potentially immunotherapy-refractory tumor microenvironments.

**Figure 7.**
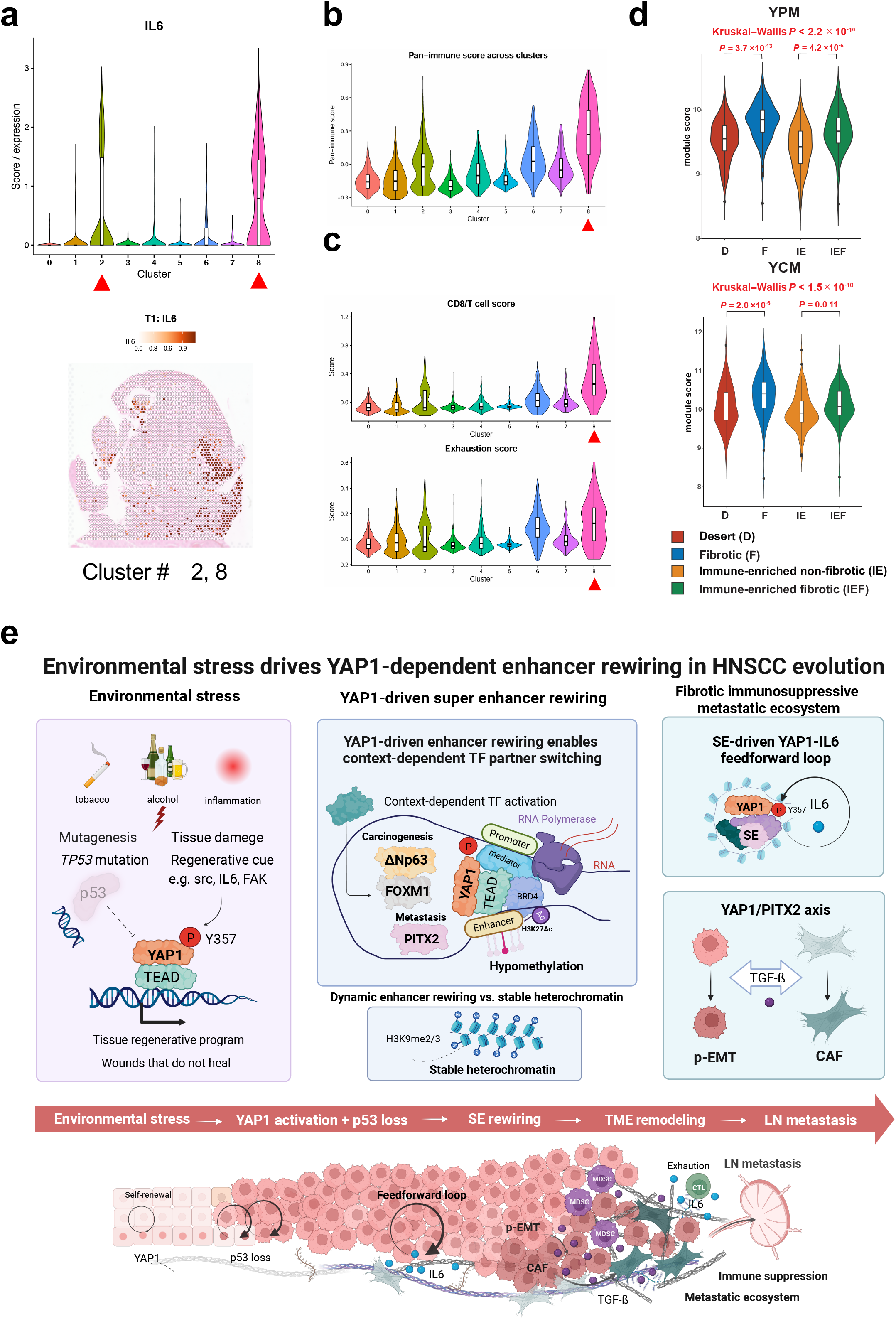
Spatially compartmentalized immune ecosystems associated with YAP1-driven metastatic progression. **a** Violin plot and spatial map showing IL6 expression across Visium ecosystems. High IL6 expression is predominantly confined to cluster 8, an immune-rich vascular stromal ecosystem located adjacent to fibrotic invasive niches. **b** Violin plot of pan-leukocyte signatures. **c** Violin plot of CD8□ T-cell signatures and exhaustion-associated transcriptional programs. CD8□ T cells are largely restricted to cluster 8 and display an exhaustion-like phenotype. **d** Association between YAP1-associated transcriptional programs and tumor immune phenotypes in the TCGA HNSCC cohort. Violin plots show YPM (upper) and YCM (lower) module scores across molecular functional tumor portrait subtypes: desert (D), fibrotic (F), immune-enriched (IE), and immune-enriched fibrotic (IE/F). Overall differences among groups were assessed using the Kruskal–Wallis test, followed by pairwise two-sided Wilcoxon rank-sum tests with Bonferroni correction. Bonferroni-adjusted P values are shown for D versus F and IE versus IE/F comparisons. **e** Graphical summary of the proposed mechanism. Environmental stressors induce tissue injury and *TP53* mutation, leading to persistent YAP1 activation. YAP1 drives super-enhancer rewiring accompanied by enhancer hypomethylation and cooperates with stage-specific transcription factors, including FOXM1, ΔNp63, and PITX2, to promote transcriptional evolution. Downstream activation of p-EMT, IL6 signaling, CAF activation, and immune remodeling establishes a fibrotic and immunosuppressive metastatic ecosystem that facilitates lymph node metastasis. Created in BioRender. https://BioRender.com/hnlub6y. See Source Data for a, b, and c.

Collectively, these findings support a model whereby environmental stress-induced YAP1 activation, in the context of p53 dysfunction, drives super-enhancer rewiring and transcriptional evolution through stage-specific transcription factor partnerships. This YAP1-centered epigenetic circuit coordinates epithelial, stromal, and immune remodeling to establish a fibrotic and immunosuppressive metastatic ecosystem that promotes lymph node metastasis in HNSCC (Fig. 7e).

## Discussion

In the era of whole-genome sequencing, mutation-oriented precision medicine has shown limited success in HNSCC ^60^. Notably, the most frequently observed genetic alteration, *TP53* mutation, is insufficient to initiate tumorigenesis in both animal models and human tissues ^5,61^. These observations suggest that genetic alterations are necessary but not sufficient to drive HNSCC development. Instead, our findings support a model in which HNSCC is driven by gene–environment–dependent transcriptional reprogramming. Within this framework, YAP1 emerges as a central regulator that integrates environmental cues into oncogenic transcriptional programs. The potent oncogenic role of YAP1 in HNSCC has been established in our previous animal study ^19^ and is further reinforced by the human multi-omics data presented here. Importantly, our results demonstrate that *TP53* mutation facilitates YAP1-driven transcriptional reprogramming, thereby establishing an accelerator–brake imbalance in which YAP1 activation is sustained due to loss of p53-mediated constraint. This concept is consistent with prior observations that p53 acts as a barrier to cellular reprogramming, whereas YAP1 promotes regenerative transcriptional states, as exemplified in induced pluripotent stem cell models^62,63^.

An important finding of this study is that global chromatin architecture remains largely preserved during HNSCC evolution, whereas profound changes occur at the level of enhancer regulation. Specifically, YAP1 dynamically redistributes to super-enhancers and promotes enhancer DNA hypomethylation, leading to activation of oncogenic transcriptional modules. These findings indicate that HNSCC progression is driven not by widespread chromatin disruption but by selective enhancer rewiring, providing a mechanistic basis for transcriptional addiction in this disease.

Beyond transcriptional rewiring, spatial transcriptomic analysis revealed that these molecular alterations are accompanied by coordinated remodeling of the tumor ecosystem. Rather than occurring uniformly throughout the tumor, YAP1-associated transcriptional programs evolved along a spatial epithelial–stromal remodeling axis extending from superficial tumor regions toward the invasive front. This transition was characterized by progressive activation of YCM, YC3M, YC5M, and ultimately YPM, together with enrichment of TGF-β-induced CAF and p-EMT signatures. Histopathological validation further demonstrated that YAP1-positive budding tumor cells were closely associated with YAP1-positive CAFs, supporting the concept of a fibrotic metastatic ecosystem. An additional insight from the spatial analysis is that immunosuppressive programs were not uniformly distributed but spatially compartmentalized. MDSC-like ecosystems were localized within tumor-associated epithelial compartments, whereas macrophage-rich and T-cell-rich ecosystems formed distinct stromal compartments. These findings suggest that YAP1-driven tumor progression is accompanied by organization of spatially distinct immune ecosystems rather than simple immune exclusion.

Therapeutically, these findings highlight YAP1-associated transcriptional machinery as a promising target in HNSCC. Recent efforts to disrupt YAP1–TEAD interactions, including inhibitors of TEAD palmitoylation, are currently under clinical investigation ^64^. In addition, epigenetic therapies targeting super-enhancer complexes, such as BET inhibitors ^65^, may be particularly effective in tumors exhibiting YAP1-driven enhancer reprogramming. Furthermore, given the feedforward interaction between YAP1 and IL6 signaling identified in this study, combined targeting of transcriptional and inflammatory pathways may represent a rational therapeutic strategy. Importantly, YAP1 inhibition is expected not only to suppress tumor cell proliferation but also to disrupt enhancer-driven transcriptional programs and tumor ecosystem remodeling, including CAF activation and inflammatory signaling. Notably, by attenuating TGF-β–driven fibrosis and inflammatory signaling, YAP1 inhibition may alleviate the immunosuppressive tumor microenvironment, thereby enhancing antitumor immunity and potentially improving responsiveness to immune checkpoint inhibitor (ICI) in HNSCC.

Collectively, our findings redefine HNSCC as an enhancer-driven disease in which environmental stress activates YAP1 while disabling the *TP53* brake, thereby promoting enhancer rewiring, spatial ecosystem remodeling, and metastatic evolution.

### Limitation

This study has several limitations. The multi-omics analyses were performed on a limited number of patient samples, and further validation in larger independent cohorts is warranted. In addition, while our data support a central role for YAP1 in enhancer reprogramming, the precise molecular mechanisms underlying enhancer DNA hypomethylation remain to be fully elucidated.

## Conclusion

In conclusion, our findings establish YAP1 as a central integrator of gene–environment interactions that links environmental stress to enhancer rewiring, transcriptional evolution, and spatial tumor ecosystem remodeling in HNSCC. By coupling regenerative signaling with *TP53* mutation-mediated loss of transcriptional constraint, persistent YAP1 activation drives epithelial–stromal remodeling, fibrotic metastatic ecosystem formation, and malignant progression. These findings redefine HNSCC as an enhancer-driven disease and provide a conceptual framework for therapeutic targeting of epigenetic and microenvironmental dependencies in cancer.

## Methods

### Cell lines and culture

Mouse i*Mob1*DKO cells, human SCC9 (YAP1 low expression) cells with a stable DOX-inducible YAP1 overexpression system and human HSC4 (YAP1 high expression) cells with a stable DOX-inducible shYAP1 knockdown system were established in our previous study^19^. Human HNSCC cell lines Cal27 (YAP1 low expression) and HSC4 were obtained from the Japanese Collection of Research Bioresources. Cells were maintained in conditions described in our previous studies.

### Antibodies

For immunohistochemistry (IHC), antibodies against YAP1 (Sigma-Aldrich, WH0010413M1, 1:100), p53 (Dako, clone DO-7, M7001, 1:1000), ΔNp63 (p40; Roche, clone BC28, ready-to-use), FOXM1 (Santa Cruz Biotechnology, sc-376471, 1:100), PITX2 (Thermo Fisher Scientific, PA5-98817, 1:100), and BRD4 (Abcam, ab84776, 1:200) were used.

For immunoprecipitation and immunoblotting, primary antibodies against YAP1 (Cell Signaling Technology, 4912, 1:1000) and PITX2 (Thermo Fisher Scientific, PA5-98817, 1:1000) were used. Immunocomplexes were detected using a goat anti-rabbit IgG–HRP-conjugated secondary antibody (Cell Signaling Technology, 7074, 1:2000). For immunofluorescence, an antibody against phospho-YAP1 (Y357) (Abcam, ab62751, 1:100) and an Alexa Fluor 488-conjugated secondary antibody (Molecular Probes, 1:500) were used. Nuclei were counterstained with 4′,6-diamidino-2-phenylindole (DAPI; Wako, 1 μg/mL).

For chromatin immunoprecipitation sequencing (ChIP-seq), antibodies against H3K9me2 (MAB Institute, MAB-1037), H3K9me3 (Active Motif, 39161), H3K27ac (MAB Institute, MAB-10309; Active Motif, 39133), and YAP1 (Novus Biologicals, NB110-58358; US Biological, Y1200-01D) were used.

### Microarray analysis

Total RNA was isolated from mouse i*Mob1*DKO cells treated with or without tamoxifen (TAM; 0.5 μM) for 3 days using an RNeasy Mini Kit (Qiagen) according to the manufacturer’s instructions. One sample was analyzed for each condition (n = 1 per condition). Following assessment of RNA quantity and quality, amplified cRNA was hybridized, scanned, and analyzed according to the manufacturer’s protocol (Agilent). Raw signal intensities were log2-transformed and quantile-normalized using the preprocessCore package in Bioconductor. Genes with Z scores > 2.0 or < −2.0 were considered descriptively upregulated or downregulated, respectively. Because only one sample per condition was analyzed, no inferential statistical testing was applied to the microarray comparison.

### Whole exome sequencing (WES) and *TP53* target sequence

WES for mouse tumors was conducted as previously described ^66^. For *TP53* target sequence in exons 4, 5, 6, 7, 8, and 9, genomic DNA was extracted from formalin-fixed, paraffin-embedded tumor tissues and direct (Sanger) sequencing were performed by LSI Medience Corporation (Tokyo, Japan). Sequence variants were identified by comparison with the *TP53* gene reference sequence (NG_017013.2), and amino acid changes were annotated based on the *TP53* reference transcript (NM_000546.6).

### Public data analysis and expression module analysis

RNA-seq and clinical data for HNSCC patients were obtained from The Cancer Genome Atlas (TCGA) through the Broad Institute Firehose portal (http://gdac.broadinstitute.org/runs/stddata__2016_01_28/data/HNSC/20160128/). The p-EMT signature, HNSCC single-cell RNA-seq dataset (GSE103322), and TGF-β-induced CAF signature were obtained from previously published studies^55,58^. Activities of YCM, YC3M, YC5M, and YPM were quantified using single-sample gene set enrichment analysis (ssGSEA) as previously described ^22^. For comparisons between normal and tumor tissues, module activities were analyzed as continuous variables. For survival analyses, optimal cutoff values were determined using maximally selected rank statistics implemented in the maxstat package, and patients were stratified into high- and low-activity groups. Single-cell RNA-seq data were analyzed using Seurat v4.3.0, and dimensionality reduction was performed using Uniform Manifold Approximation and Projection (UMAP). Tumor microenvironment (TME) subtypes were assigned using the Molecular Functional Tumor Portrait platform (https://science.bostongene.com/tumor-portrait/) ^59^.

### Ethics Statement

The protocol of this study was reviewed and approved by the institutional review boards and ethics committees of National Kyushu Cancer Center Kyushu University (2019–024). All experiments with human samples were conducted according to the principles expressed in the Declaration of Helsinki.

### Patients and sample collection

FFPE and frozen HNSCC tumor and adjacent normal tissue samples were collected from treatment-naïve patients who underwent radical surgery at National Kyushu Cancer Center, Department of Head and Neck Surgery. Written informed consent was obtained from all patients.

### RNA sequencing and bioinformatics analysis

Total RNA was isolated from human HNSCC cell lines and samples using a NucleoSpin RNA Plus kit (Macherey-Nagel, Germany). RNA sequencing was performed on a DNBSEQ-G400 sequencer at Beijing Genomics Institutions (Shenzhen, China) as previously described. The sequencing reads were aligned to the human reference GRCh38/hg38 genome by STAR v2.7.9 using GENCODE v38 annotations. Gene count tables were generated using RSEM v1.3.1. Downstream analyses were carried out using R v4.4.1 (R Foundation for Statistical Computing). Read count normalization and differential expression analysis were performed using DESeq2 v1.10.1. Genes with zero counts across all samples were excluded before principal component and sample -clustering analyses. Over-representation analysis of differentially expressed genes was performed using Hallmark gene sets, Gene Ontology Biological Process terms, and Kyoto Encyclopedia of Genes and Genomes (KEGG) pathways with the enrichR v3.2. Enrichment results were visualized as dot plots, with the gene ratio or numbers shown on the x axis, dot size representing the number of overlapping genes, and color indicating the Benjamini–Hochberg-adjusted P value. Conventional gene set enrichment analysis was performed using GSEA v4.3.2 on ranked gene lists. Enrichment was evaluated using the normalized enrichment score (NES) and false discovery rate (FDR) q value. Gene signature activities in individual samples were quantified by single-sample gene set enrichment (ssGSEA) analysis implemented in GSVA v1.52.3. For survival analyses, optimal cutoff values were determined using maximally selected rank statistics implemented in the maxstat v0.7-25, after which patients were stratified into high- and low-score groups. Specific gene set–based transcriptome analysis was performed using the Expression Enrichment Matrix (EEM), a framework that converts gene expression data into gene set–level enrichment scores across samples ^56^. Gene sets used for EEM analysis included curated gene sets from the Molecular Signatures Database (MSigDB), together with study-defined gene sets derived from YAP1-dependent differentially expressed genes identified by RNA-seq following YAP1 perturbation in HSC4 and SCC9 cells and genes associated with YAP1-regulated super-enhancers. These gene sets encompassed functional programs related to cell cycle, inflammation, EMT, matrix remodeling, YAP1-associated transcriptional programs, TGF-β signaling, and immune responses. Study-defined gene sets and their constituent genes are provided in Supplementary Data 3.TF activity prediction was performed using R package decoupleR v2.9.1 ^67^.

### ChIP sequencing

ChIP-seq experiments using mouse-derived cells and human HNSCC cell lines were performed at the Laboratory of Genome Structure and Function, Institute for Quantitative Biosciences, The University of Tokyo, with support from the Platform Project for Supporting Drug Discovery and Life Science Research (BINDS) of AMED (grant no. 0667), using a previously described protocol ^68^. Chromatin immunoprecipitation was performed using antibodies against H3K9me2 (MAB Institute, MAB-1037), H3K9me3 (Active Motif, 39161), H3K27ac (MAB Institute, MAB-10309; Active Motif, 39133), and YAP1 (Novus Biologicals, NB110-58358; US Biological, Y1200-01D), as appropriate for each experiment. ChIP-seq libraries were prepared using the NEBNext Ultra II DNA Library Prep Kit for Illumina (New England Biolabs, E7645) according to the manufacturer’s instructions, except that size selection was performed after PCR amplification using AMPure XP beads (Beckman Coulter, A63881). Libraries were sequenced on an Illumina HiSeq 2500 platform to generate 65-bp single-end reads.

Frozen HNSCC tumor samples were sent to Active Motif (Carlsbad, CA, USA) for chromatin immunoprecipitation, library preparation, and sequencing as previously described ^22^. In brief, chromatin was immunoprecipitated using antibodies against H3K27ac (Active Motif, 39133), H3K9me3 (Active Motif, 39161), and YAP1 (US Biological, Y1200-01D,). ChIP-seq libraries were sequenced on an Illumina NextSeq 500 platform to generate 75-bp single-end reads.

### ChIP-seq data analysis

Mouse ChIP-seq reads were aligned to the mm10 reference genome using Bowtie. Wiggle files were generated using parse2wig in DROMPA3 v3.2.3^69^ and d DROMPA3 was used for visualization and peak calling. Peaks were called in PC_SHARP mode using the corresponding input sample as control and default parameters. Genome-wide ChIP/Input enrichment profiles were visualized using the GV mode, and normalized profiles were scaled to 20 million mapped reads for comparative analyses. The same FASTQ datasets were additionally analyzed using MACS2 v2.1.0 for peak calling in the analyses shown in Supplementary Fig. 2b, with a *P*-value cutoff of 1 × 10□□ and otherwise default parameters. Peak-centered signal profiles and heatmaps were generated using deepTools, and the genomic distribution of called peaks was annotated and visualized using the R package ChIPseeker..

Human HNSCC cell-line ChIP-seq reads were aligned to the human reference genome GRCh38/hg38. Peaks were called using MACS2 v2.1.0 with a *P*-value cutoff of 1 × 10□□ and otherwise default parameters, and EPIC was used for broad enrichment analysis. ChIP-seq signal profiles and heatmaps were generated using deepTools, and genomic annotation of peaks was performed using the R package ChIPseeker.

For human HNSCC specimens, 75bp single-end raw reads were aligned to the human reference genome GRCh38/hg38 using BWA v0.7.12. Peaks were identified using MACS2 v2.1.0 with a *P*-value cutoff of 1 × 10□□. ChIP-seq signal profiles and heatmaps were generated using deepTools v3.5.5, and genomic annotation of called peaks was performed using the R package ChIPseeker. For multiregional HNSCC samples, ChIP-seq enrichment profiles were clustered into five clusters (*k* = 5) using deepTools. SEs were identified using the ROSE algorithm with default parameters ^70^. Overlapping SE regions were determined using multiinter in BEDTools v2.31.1, and regions shorter than 3 kb were excluded. Enhancer-to-gene interactions in the detected enhancers and SEs were inferred using GeneHancer v5.24, a comprehensive enhancer/promoter database of human genome ^71^, after excluding regions annotated as promoters. Nucleosome-free regions (NFRs) within H3K27ac-enriched regions were inferred using the HisTrader algorithm as previously described ^72^. YAP peaks were annotated to putative transcriptional target genes using GREAT ^73^ with the Basal plus Extension option (2kb upstream and 2kb downstream, distal up to 1,000kb options).

ChIP-seq signal profiles, heatmaps, and genomic regions were visualized using deepTools v3.5.5, ChIPseeker, and pyGenomeTracks v3.9, as appropriate. Representative genome-browser views, including the *IL6* and *PDPN* locus, were generated using Integrative Genomics Viewer (IGV) v.2.19.1.

### Whole genome bisulfite sequencing (WGBS) and reduced representation bisulfite sequencing (RRBS)

WGBS was performed on mouse-derived control and i*Mob1*DKO cells collected at 1 and 2 weeks after TAM treatment at the Department of Biochemistry, Kyushu University Graduate School of Medical Sciences, with support from the Platform Project for Supporting Drug Discovery and Life Science Research (BINDS) from AMED (grant no. 0668). WGBS libraries were prepared using the post-bisulfite adaptor tagging (PBAT) strategy ^74^, following the improved protocol incorporating highly efficient single-stranded DNA ligation ^75^. The molar concentration of the sequencing libraries was determined using a Library Quantification Kit (Takara Bio Inc., Shiga, Japan). Paired-end sequencing (2 × 150 bp) was performed on a HiSeq X Ten platform by Macrogen Japan Corp. (Tokyo, Japan). Sequencing reads were mapped to the mouse reference genome NCBI37/mm9 using BMap, as described in the original PBAT study^74^. Differentially methylated regions (DMRs) between samples were identified using the DSS R package with a nominal P-value cutoff of 0.01.

Human HNSCC samples for RRBS were sent to Active Motif (Carlsbad, CA, USA) for RRBS library preparation and sequencing. Sequencing reads were processed and aligned to the human reference genome GRCh38 (hg38) using a bisulfite sequencing analysis pipeline. CpG methylation levels were extracted and converted to beta values for downstream analyses. YAP1-associated super-enhancer (SE) regions were defined by integrating H3K27ac ChIP-seq, YAP1 ChIP-seq, and nucleosome-free region (NFR) estimation analyses. Cluster 3- and cluster 5-associated YAP1 SE regions were identified using merged NFR-containing enhancer regions. CpG methylation profiles surrounding C3 and C5 YAP1-binding peaks were aggregated across ±2-kb regions centered on YAP1 peak summits using 50-bp bins. For comparisons between SEs harboring C3 or C5 YAP1-binding peaks and background SEs, mean CpG methylation levels were calculated within ±500 bp of the corresponding YAP1 peak centers. Statistical significance was assessed using the Wilcoxon rank-sum test or Kruskal–Wallis test followed by Dunn’s multiple-comparison test, as appropriate.

### MOTIF analysis

Transcription factor binding motif enrichment analysis in 200bp regions around YAP peak summits and motif enrichment analysis in NFRs were performed using HOMER v5.1 ^76^ with default database included in HOMER and HOCOMOCO v13 database ^77^and significance was determined based on Benjamini–Hochberg adjusted q-values (q<0.05). De novo motif discovery was conducted using MEME(https://meme-suite.org/meme) ^78^, and motifs with E-values <1×10□□ were considered significant.

### Co-immunoprecipitation (Co-IP) assay

Co-IP was performed using SureBeads Protein A/G magnetic beads (Bio-Rad Laboratories) according to the manufacturer’s protocol. Briefly, HSC-4 cells at 70% confluency were lysed with Pierce IP Lysis Buffer (Thermo Fisher Scientific) containing protease inhibitors on ice, and protein concentration was determined using a BCA protein assay kit (Takara Bio). Total cell lysate (0.5 mg) was incubated with YAP antibody (1:1000, #4912, Cell Signaling Technology) or control IgG for 2 h at room temperature, followed by incubation with magnetic beads at 4°C overnight. The immunoprecipitates were washed three times with PBS containing 0.1% Tween 20 (PBS-T). Bound proteins were eluted with 1x SDS sample buffer (Bio-Rad Laboratories) at 98°C for 5 min. Eluted samples along with input samples were loaded onto 10% Mini-PROTEAN TGX gels (Bio-Rad Laboratories), separated by SDS-PAGE, and transferred onto PVDF membranes. After blocking with 5% non-fat milk in TBS-T, membranes were incubated overnight at 4°C with primary antibodies: PITX2 (1:1000, #PA5-98817, Thermo Fisher Scientific) and YAP (1:1000, #4912, Cell Signaling Technology). Membranes were then incubated with goat anti-rabbit IgG/HRP-conjugated secondary antibody (1:2000, #7074, Cell Signaling Technology) for 1 h at room temperature. Protein bands were visualized using Clarity ECL Western Blotting Substrate and the ChemiDoc MP Imaging System (Bio-Rad Laboratories).

### siRNA-mediated knockdown experiments

HSC4 cells were seeded in 6-well plates at 2 × 10□ cells per well and incubated overnight. Two independent siRNAs targeting PITX2 (s10558 and s10559; Thermo Fisher Scientific) and two independent siRNAs targeting IL6 (s7311 and s7312; Thermo Fisher Scientific) were used together with Silencer Select Negative Control No. 1 siRNA (Thermo Fisher Scientific). siRNAs were transfected at a final concentration of 20 nM using Lipofectamine RNAiMAX (Thermo Fisher Scientific) according to the manufacturer’s protocol. Cells were harvested 48 h after transfection for RNA or protein extraction and were also used for downstream functional assays.

The siRNA sequences were as follows:

si-PITX2 #1, Sense; ACUCCUCCGUAUGUUUAUAtt, Antisense;

UAUAAACAUACGGAGGAGUcg

si-PITX2 #2, Sense; GGACAUGUCCACACGCGAAtt, Antisense;

UUCGCGUGUGGACAUGUCCgg

si-IL6 #1, Sense; GAACGAAUUGACAAACAAAtt, Antisense;

UUUGUUUGUCAAUUCGUUCtg

si-IL6 #2, Sense; CUUCCAAUCUGGAUUCAAUtt, Antisense;

AUUGAAUCCAGAUUGGAAGca

### Sphere formation assay

Cells were seeded onto 6-well ultra-low attachment plates (Corning) at a density of 5,000 cells/well in Ham’s F12 containing 20 ng/ml EGF (#AF-100-15, PeproTech, USA), 20 ng/ml bFGF (#100-18B, PeproTech) and B27 Supplement (#17504044, Invitrogen). Triplicate wells were prepared for each cell. After 2 weeks, visible spheres were manually counted using a microscope.

### Histological analyses and Immunostaining

The protein expression levels of YAP1, p53, FOXM1, deltaNp63(p40), BRD4, and PITX2 in the primary HNSCC tissues were evaluated by immunohistochemistry (IHC) as previously described ^19,20,22^. Briefly, nuclear staining intensity was defined as “intensity score” (scored as 0–3), whereas the population of positive-staining cells was defined as “frequency score” (scored as 0–4). YAP1expression scores were defined by the following formula: (intensity score) × (frequency score), and samples with a score > 7 were classified as the YAP1-high group. For BRD4 and PITX2 samples with a intensity score >2 were classified as the high group. The histology and the results of IHC were independently reviewed by three experienced pathologists (T.H., T.N., and T.K.). Immunofluorescence staining for HNSCC cells was performed as previously described ^19^ using an indirect method. Cells were stained for phospho-YAP1 (Y357), followed by detection with an Alexa Fluor 488-conjugated secondary antibody. Nuclei were counterstained with DAPI, and samples were mounted using PermaFluor (Thermo Scientific). Fluorescence images were acquired using a Leica SP8 STED confocal microscope.

### Spatial transcriptomics (Visium)

Spatial transcriptomic analysis was performed using the Visium Spatial Gene Expression for FFPE platform (10x Genomics) on multiregional HNSCC specimens corresponding to those used for multi-omics analyses. Regions of interest (ROIs) exhibiting high nuclear YAP1 expression were identified by immunohistochemistry and subjected to spatial transcriptomic profiling. Library preparation and sequencing were performed according to the manufacturer’s protocols. Raw sequencing data were processed using Space Ranger v2.0.1 (10x Genomics) and aligned to the human reference genome GRCh38 (hg38). Downstream analyses were performed using Seurat v4.3.0 in R v4.4.1. Dimensionality reduction and clustering were performed using principal component analysis (PCA) and UMAP. Clusters were annotated based on canonical marker genes, spatial localization patterns, and inferred cellular compositions obtained using cell2location v0.1.4. Gene set activity scores, including YAP1-associated modules (YCM, YC3M, YC5M, and YPM), p-EMT signatures, CAF signatures, and TGF-β-induced CAF signatures, were calculated using the AddModuleScore function implemented in Seurat. Spatial visualization of gene expression and module activities was performed using SpatialFeaturePlot. Gene signatures and marker gene sets used for functional and spatial annotation are provided in Supplementary Data 3.

### Statistical Analysis

Statistical analyses were performed using R v4.4.1 or JMP software (version 19.0.1). Overall survival and progression-free survival were estimated using the Kaplan–Meier method and compared using the log-rank test. Hazard ratios and 95% confidence intervals were estimated using univariable and multivariable Cox proportional hazards models. Categorical variables were compared using Fisher’s exact test. Continuous variables were compared using a two-sided unpaired t-test, Mann–Whitney U test, or Kruskal–Wallis test, as appropriate. For comparisons among three or more groups, post hoc multiple-comparison testing was performed using either Dunn’s test following the Kruskal–Wallis test or pairwise two-sided Wilcoxon rank-sum tests with Bonferroni correction, as indicated. For analyses of YAP1-associated module scores across Molecular Functional Tumor Portrait subtypes, overall differences among the four groups were assessed using the Kruskal–Wallis test, followed by pairwise two-sided Wilcoxon rank-sum tests with Bonferroni correction; Bonferroni-adjusted P values are reported. Odds ratios (ORs) were calculated, and when zero counts were present, the Haldane–Anscombe correction (addition of 0.5 to all cells) was applied to estimate corrected ORs and 95% confidence intervals. Data are presented as mean ± SEM unless otherwise indicated. P values < 0.05 were considered statistically significant.

## Data Availability

Mouse tumor whole-exome sequencing data have been deposited in the NCBI Sequence Read Archive (SRA) under BioProject accession number PRJNA1558269. The RNA-seq, ChIP-seq, RRBS, WGBS, and microarray data generated in this study have been deposited in the NCBI Gene Expression Omnibus (GEO) database under the following Series accession numbers: GSE353915 (human cell line RNA-seq), GSE353917 (human tissue ChIP-seq), GSE353918 (human tissue RNA-seq), GSE353919 (human tissue RRBS), GSE353920 (mouse ChIP-seq), GSE353924 (human cell line ChIP-seq), GSE353941 (mouse WGBS), and GSE353944 (mouse microarray). The spatial transcriptomics (Visium) data are available in GEO under accession number GSE348297.

## Acknowledgement

This research was partially supported by Platform Project for Supporting Drug Discovery and Life Science Research (Basis for Supporting Innovative Drug Discovery and Life Science Research: BINDS) from AMED under 0667 and 0668.

## Author contributions

M.H., Kuniaki Sato, and H.O. contributed equally to this work. M.H., Kuniaki Sato, and H.O. conducted the experimental studies, performed bioinformatic analyses, analyzed and interpreted the data, and contributed to the preparation of figures and the manuscript. T.H., K.T., and Takafumi Nakano performed and analyzed the immunohistochemical studies. M.B., R.N., and Katsuhiko Shirahige performed the mouse ChIP-seq experiments and contributed to the analysis and interpretation of the epigenomic data. T.I., F.M., and H.A. contributed to the mouse whole-genome bisulfite sequencing experiments, WGBS data generation and curation, and DNA methylation analyses. N.K., Takuya Nakagawa, A.K., and J.S.G. supervised the research and provided scientific and technical guidance. M.M. conceived the study, developed the overall research concept, acquired funding, designed and supervised the project, interpreted the data, and wrote the manuscript. All authors reviewed and approved the final manuscript.

## SUPPLEMENTARY INFORMATION

**Supplementary Fig. 1.**
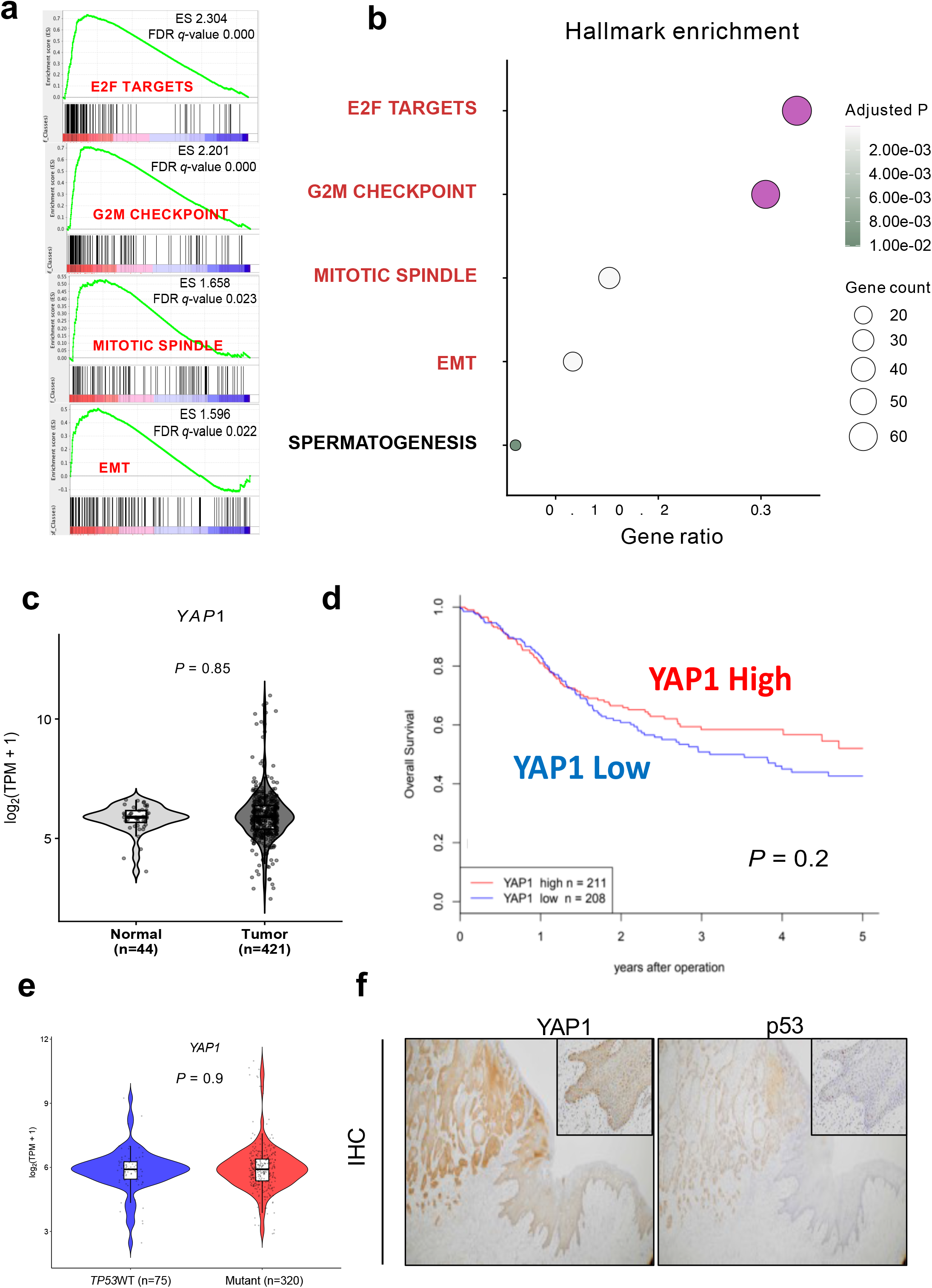
Gene set enrichment analysis, YAP1 mRNA in TCGA dataset and representative p53 and YAP1 immunohistochemical staining. **a** Representative gene set enrichment analysis (GSEA) of upregulated genes (Z score > 2) and gene ratio at 1 week after TAM treatment in *iMob1*DKOcells, corresponding to the dysplastic stage. **b** Functional annotation (Hallmarks) of upregulated genes (Z score > 2) and gene ratio at 1 week after TAM treatment in *iMob1*DKOcells, corresponding to the dysplastic stage. *P* values were calculated using a two-sided Wilcoxon rank-sum test. **C** Violin plot comparison of *YAP1* mRNA expression levels between normal and tumor samples in TCGA dataset. **d** Kaplan–Meier analysis showing no significant association in TCGA dataset between YAP1 mRNA expression and patient survival (*P* = 0.2, log-rank test). **e** Association between *TP53* mutation and *YAP1* mRNA in TCGA dataset. **f** Representative immunohistochemical staining of human HNSCC samples demonstrating colocalization of mutant p53 and YAP1 proteins in cancer-adjacent normal epithelium. See Source Data for c and e.

**Supplementary Figure 2.**
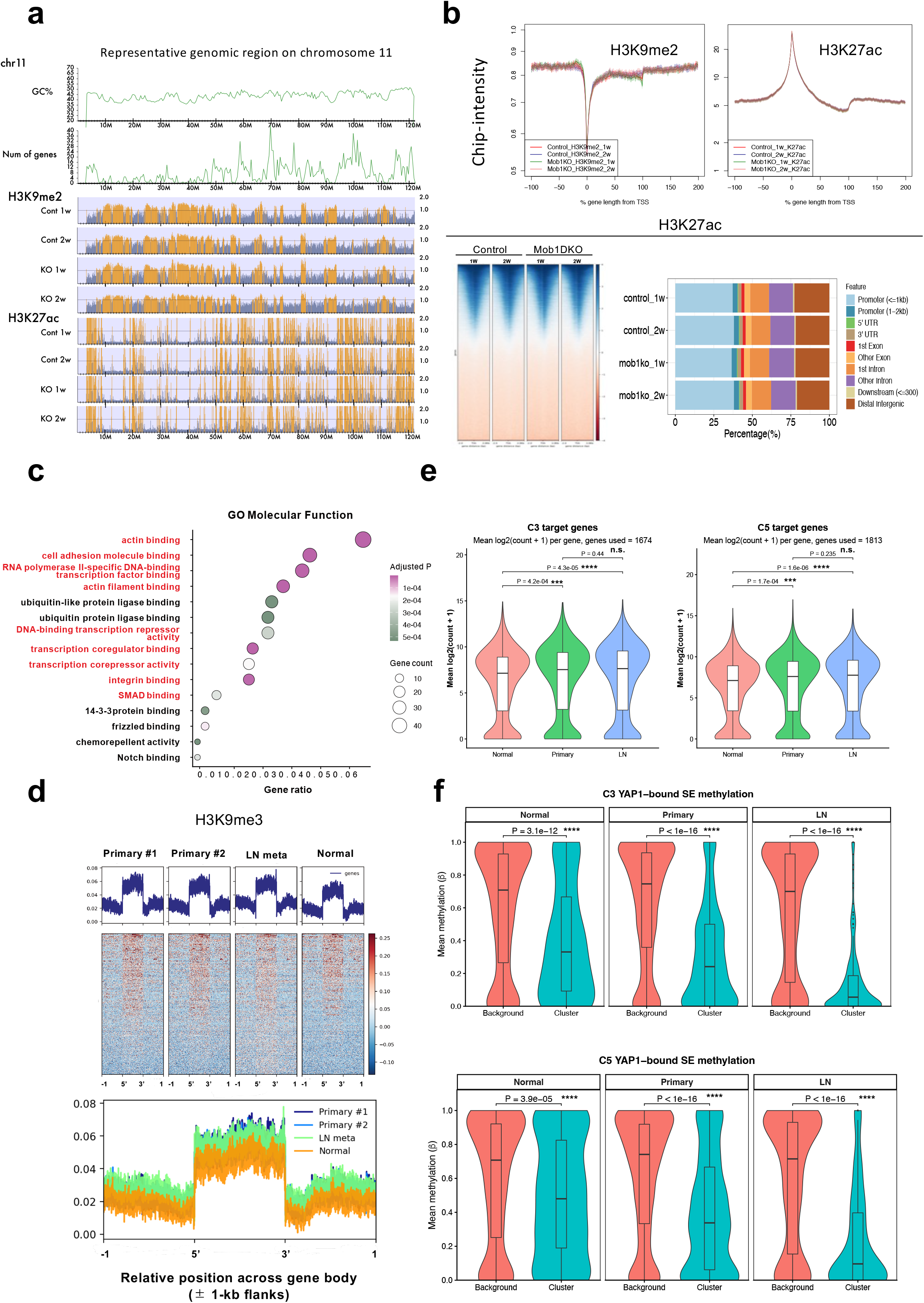
Chromatin architecture. **a** Genome browser view of representative chromosomal region (chr11) showing GC content, gene density, and ChIP-seq tracks for H3K9me2 and H3K27ac in control and *iMob1*DKO cells at 1 and 2 weeks after tamoxifen treatment. **b** Aggregate plots, heatmaps, and genomic distribution plots of H3K9me2 and H3K27ac signals across transcription start site (TSS)-proximal regions following YAP1 activation. **c** Functional annotation (GO Molecular Function) of genes associated with hypomethylated super-enhancers. The x-axis represents the gene ratio, circle size indicates gene count, and color indicates the Benjamini–Hochberg-adjusted *P.* **d** Metagene analysis of H3K9me3 ChIP-seq signals across gene bodies and flanking regions (±1kb) in normal tissue, primary tumors, and lymph node m etastases. **e** Violin plots showing the expression of YAP1 target genes associated with cluster 3 (upper) and cluster 5 (lower) in normal mucosa, primary tumors, and matched lymph node (LN) metastases. Expression of both gene sets was significantly increased in primary tumors and LN metastases compared with normal mucosa, whereas no significant difference was observed between primary tumors and LN metastases. *P* values were calculated using the two-sided Wilcoxon rank-sum test. *** *P* < 0.001; ****P < 0.0001; n.s., not significant. **f** Violin plots comparing mean DNA methylation levels within ±500 bp surrounding the centers of cluster 3 (upper) and cluster 5 (lower) YAP1-binding peaks in super-enhancers harboring these peaks with background super-enhancers in normal mucosa, primary tumors, and matched LN metastases. Super-enhancers harboring C3 or C5 YAP1-binding peaks exhibited significantly lower DNA methylation than background super-enhancers in all three tissue types, with progressive hypomethylation from normal mucosa to primary tumors and further to LN metastases. *P* values were calculated using the two-sided Wilcoxon rank-sum test ****P < 0.0001 See Source Data for e and f.

**Supplementary Fig. 3.**
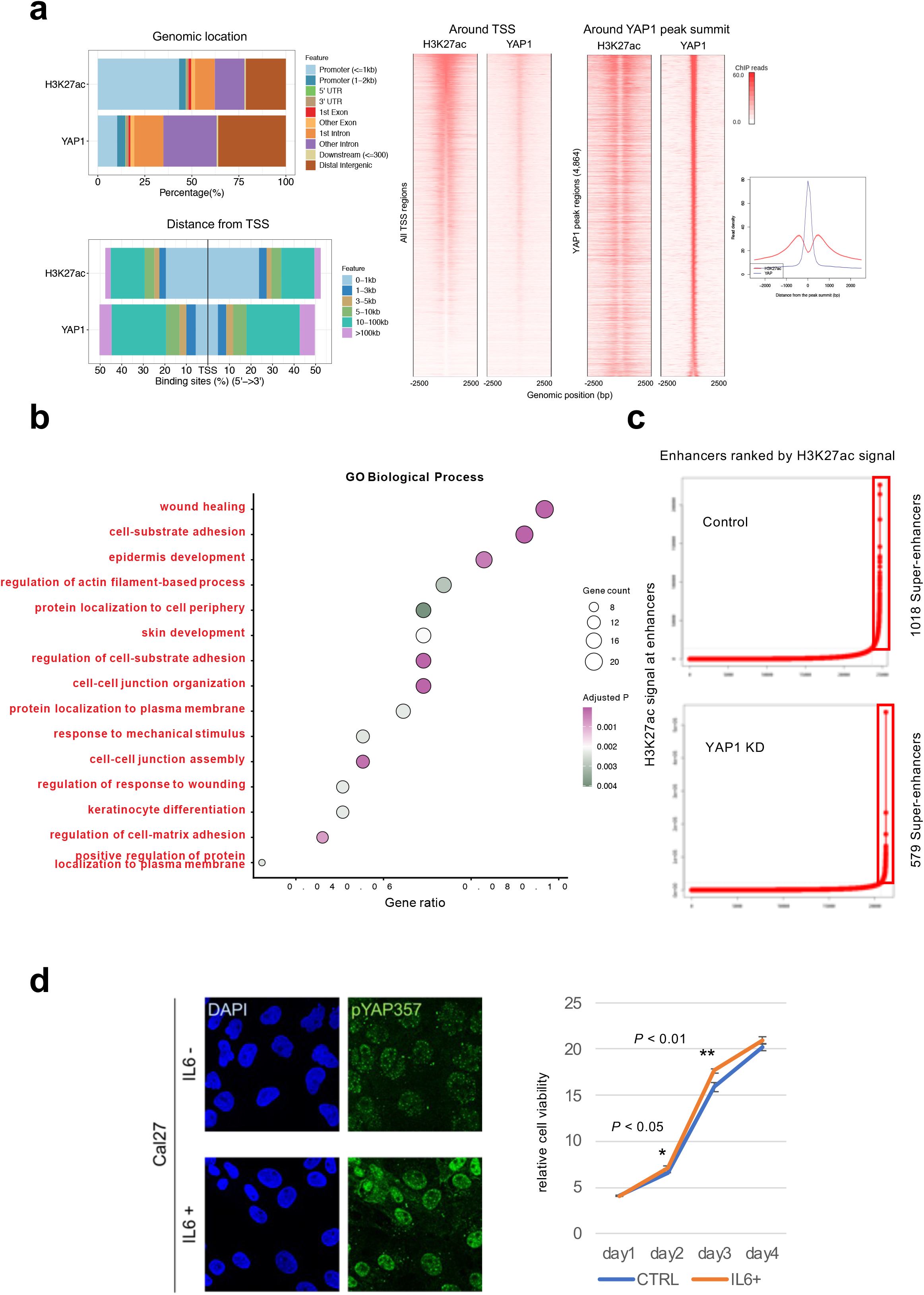
Genome-wide enhancer binding profile of YAP1 and functional validation of the YAP1–IL6 axis. **a** Genomic distribution of H3K27ac and YAP1 ChIP–seq peaks in YAP1-high HSC4 cells. **b** Functional annotation (GO Biological Process) of YAP1-bound enhancer-associated genes. The x-axis represents the gene ratio, circle size indicates gene count, and color indicates the Benjamini–Hochberg-adjusted *P* value. **c** In YAP1-low Cal27 cells, exogenous IL6 increased YAP1 Y357 phosphorylation and promoted cell growth. Data are presented as mean ± SD. Comparisons between control and IL6 were performed using two-sided Student’s t-tests at each time point. *P* < 0.05 (day 2) and P < 0.01 (day 3). See Source Data for c.

**Supplementary Fig. 4.**
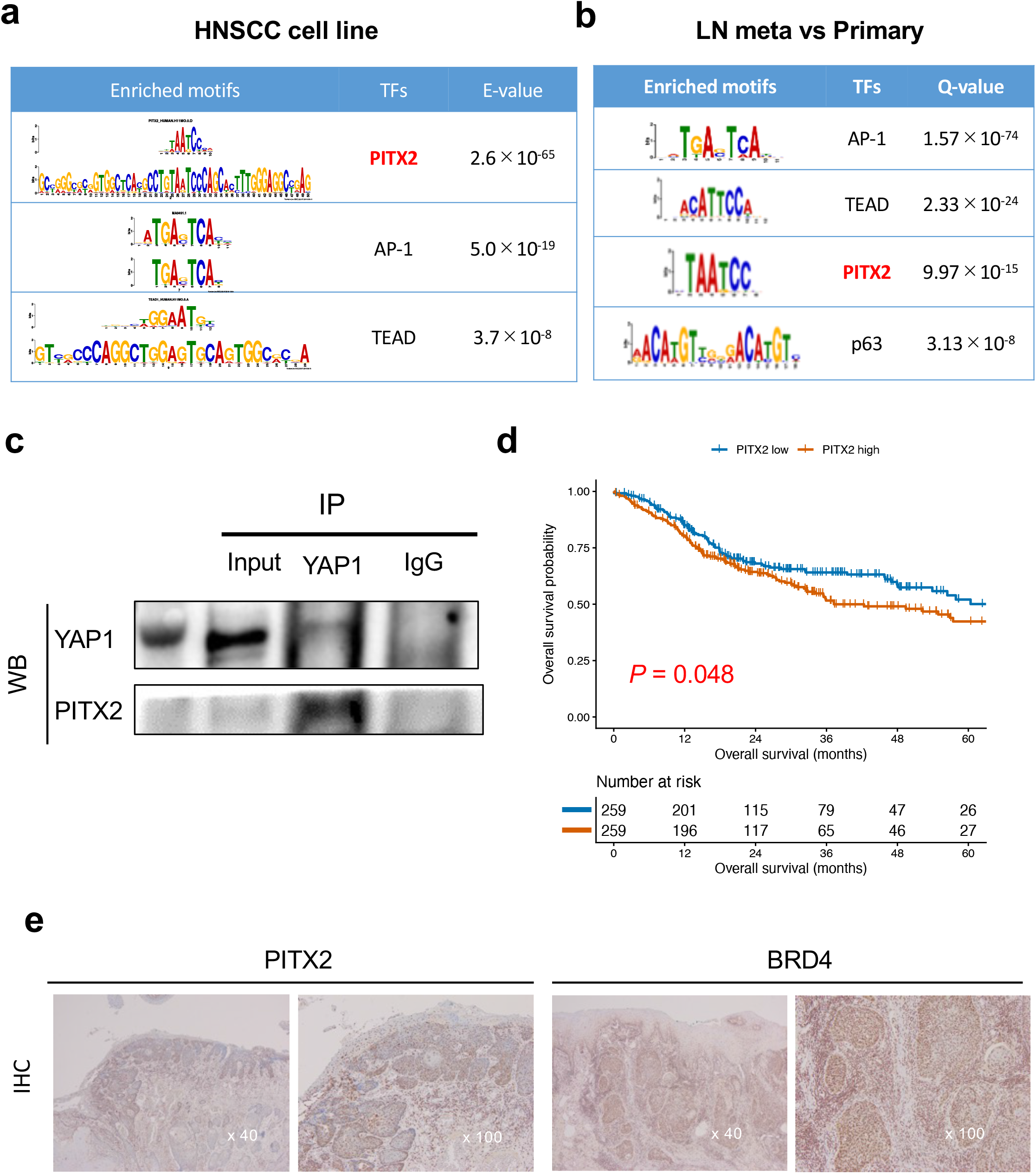
Validation of PITX2 as a progression-associated YAP1 partner transcription factor. **a** Motif enrichment analysis of YAP1-bound SE regions in HNSCC cell lines. **b** Motif enrichment analysis of genes significantly upregulated in lymph node metastases compared with paired primary tumors (RNA-seq, n = 10), identifying PITX2-binding motifs among the top enriched sequences. **c** Co-immunoprecipitation assays demonstrating physical interaction between YAP1 and PITX2 in HNSCC cell lines. **d** Kaplan–Meier analysis of overall survival in the TCGA HNSCC cohort stratified by PITX2 mRNA expression (median cutoff). High PITX2 expression was associated with poorer survival (P = 0.048, log-rank test). **e** Representative PITX2, and BRD4 positive immunohistochemical staining.

**Supplementary Fig. 5.**
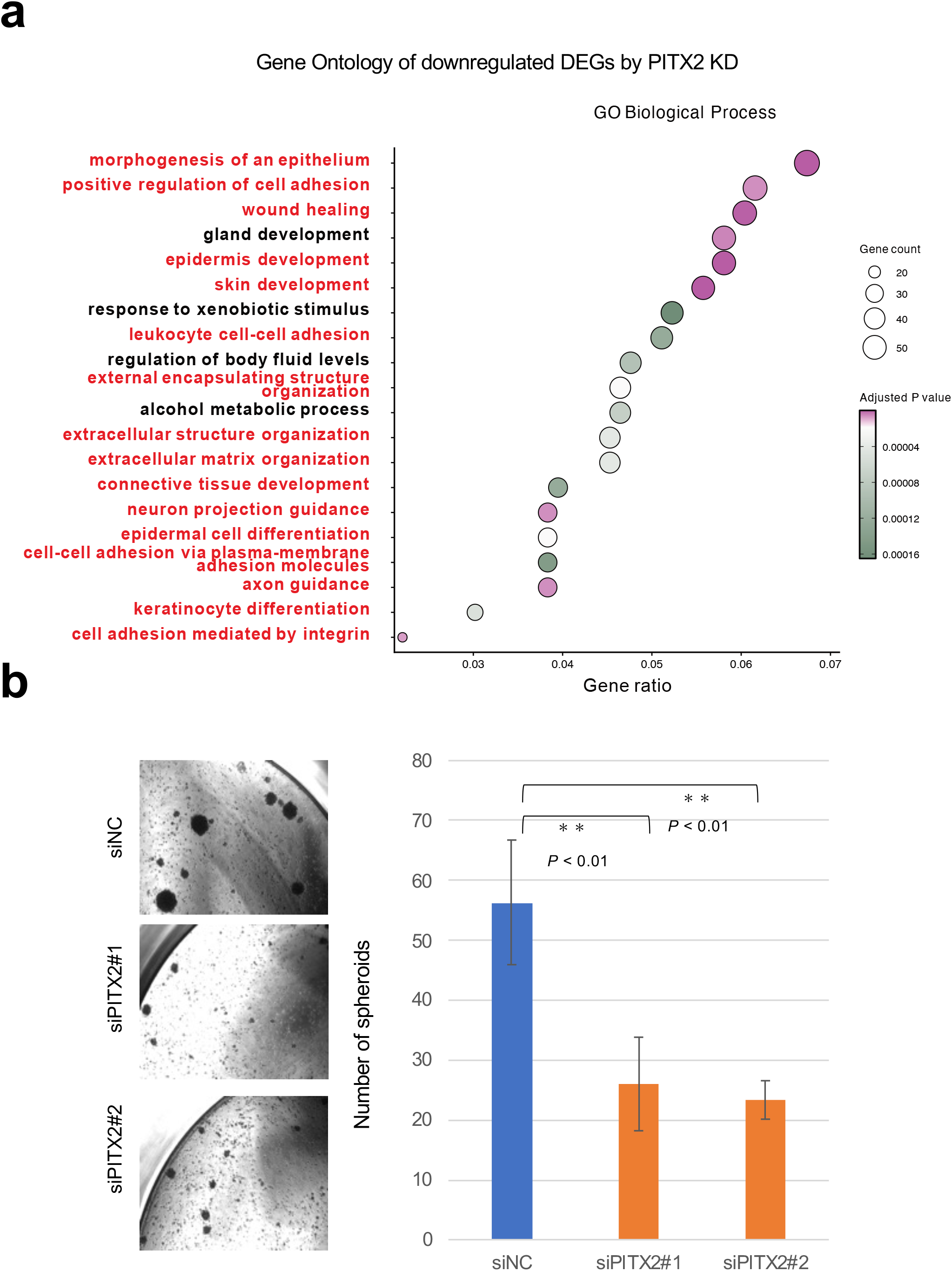
Functional consequences of PITX2 depletion in HNSCC cells. **a** Functional annotation (GO Biological Process) of downregulated DEGs following PITX2 knockdow. The x-axis represents the gene ratio, circle size indicates gene count, and color indicates the Benjamini–Hochberg-adjusted P. **b** Sphere-formation assays demonstrating reduced stemness capacity upon PITX2 silencing. Data are presented as mean ± s.d. from independent experiments. P values were calculated using the two-sided Wilcoxon rank-sum test. \*\**P* < 0.01; \*\*\**P* < 0.001.). See Source Data for c.

**Supplementary Fig. 6.**
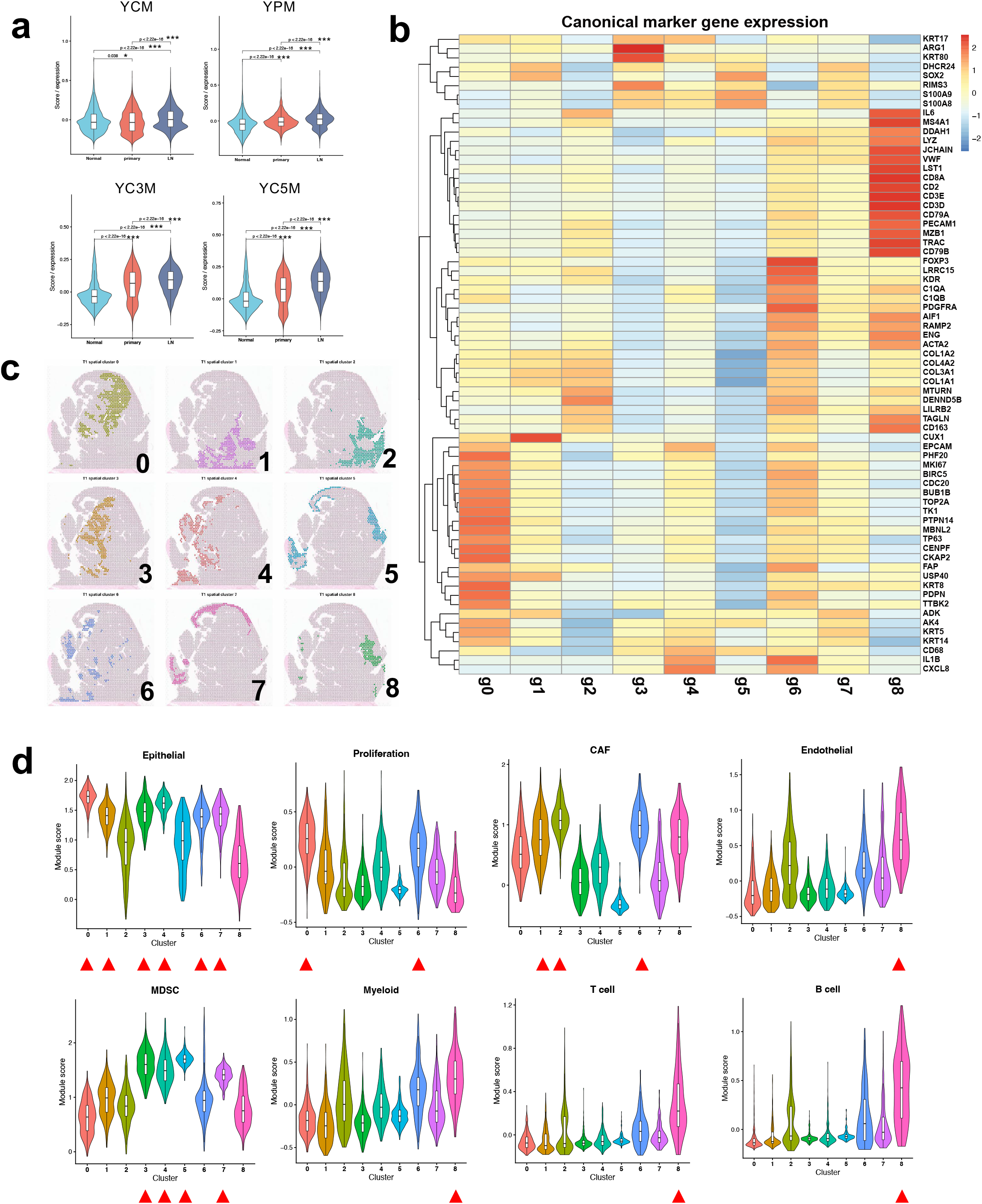
Validation of ROI selection and ecosystem annotation in spatial transcriptomic analyses. **a** Module score analysis of YCM, YPM, YC3M, and YC5M across normal epithelium, primary tumors, and matched lymph node metastases. Progressive activation of YAP1-associated transcriptional programs is observed during tumor evolution. P values were calculated using the two-sided Wilcoxon rank-sum test. \**P* < 0.05; \*\**P* < 0.01 ; ***P < 0.001. **b** Heat map of canonical lineage markers in nine clusters. **c** Spatial maps showing nine clusters. **d** Expression of representative lineage-specific marker signatures across spatial transcriptomic clusters. Violin plots showing epithelial, proliferation, cancer-associated fibroblast (CAF), endothelial, myeloid-derived suppressor cell (MDSC), myeloid, T-cell, and B-cell marker signatures across the nine clusters. These marker signatures supported the annotation of the major cellular ecosystems shown in Fig. 6c.). See Source Data for a and d.

## SUPPLEMENTARY DATA LEGENDS

**Supplementary Data 1.** Normalized microarray expression matrix used for differential expression and gene set enrichment analyses. This file contains normalized expression values for all probes detected in tongue epithelial samples from control and Mob1 double-knockout (DKO) mice at 1 and 2 weeks after YAP1 activation. These data were used for differential expression analysis and subsequent gene set enrichment analyses. **Related to Fig. 1. and Supplementary Fig. 1.**

**Supplementary Data 2.** YC3M and YC5M gene modules. **Related to Fig. 2**.

**Supplementary Data 3** Gene signatures and marker gene sets used for transcriptomic and spatial analyses。 **Related to Fig. 5,6, and 7. and Supplementary Fig. 6.**

